# Effective porosity and fluid flow in macroporous ultrasoft hydrogels: An experimental characterization

**DOI:** 10.64898/2026.04.30.721851

**Authors:** Manuel P. Kainz, Michele Terzano, Dagmar Kolb, Gerhard A. Holzapfel

**Affiliations:** Institute of Biomechanics, Graz University of Technology, Austria; Center for Medical Research, Gottfried Schatz Research Center, Core Facility Ultrastructure Analysis, Medical University of Graz, Austria; Division of Cell Biology, Histology and Embryology, Gottfried Schatz Research Center, Medical University of Graz, Austria; Department of Structural Engineering, Norwegian University of Science and Technology (NTNU) Trondheim, Norway

**Keywords:** Soft biomaterials, hydrogels, porosity, free water, porous media

## Abstract

Hydrogels are the preferred materials for applications mimicking soft tissues due to their high water content and tunable mechanical properties. The state of the water in these hydrated networks governs their response to mechanical loading through coupled interstitial flow and large deformations of the solid network. Reliable experimental methods for quantifying the fraction of mobile fluid during mechanical deformation remain limited. Within the theoretical framework of mixture theory, we describe hydrogels as hydrated biphasic media consisting of a deformable incompressible solid matrix and a mobile fluid phase. We developed a mechanical testing protocol that enables the experimental separation of solid and fluid contributions under loading. The method is demonstrated using biocompatible and highly versatile hydrogel phantoms of varying compositions. Controlled, incremental drained confined compression of the hydrogel samples results in free-water fractions of approximately 40%, 60%, and 77%, reflecting the systematic influence of the polymer content on the porosity and fluid mobility. Comparison with cryo-SEM-derived surface porosity reveals statistically significant differences and highlights the scale-dependent sensitivity of surface measurements compared to bulk measurements. This study introduces a new mechanical method for quantifying the free-water fraction in macroporous, ultrasoft, highly hydrated biomaterials. Furthermore, the multi-step protocols enable the separation of dissipative, fluid-related relaxation from the equilibrium response of the solid skeleton, allowing direct calibration of constitutive models for macroporous soft solids. The proposed method provides a reliable basis for the development and optimization of hydrogels for applications where fluid transport is critical, such as neural interfaces, bioelectronic platforms, and tissue-engineered constructs.

## 1 Introduction

Hydrogels, widely used as soft tissue phantoms [1, 2], soft actuators [3, 4], biomimetic interfaces [5, 6], and substrates or coatings for implantable devices [7–9], are polymer networks highly swollen with water, often between ten and several thousand times their dry weight [10, 11]. Transport and mechanical properties of these soft compounds are governed not only by the total water content but also by the physical state of that water [12, 13], particularly the fraction that can be mobilized under mechanical load [14]. Understanding the different states of water in hydrogels, as well as their relationship to network-architecture metrics such as porosity, is essential for designing soft, functional biomaterials with controlled mechano-responsive behavior [15–17].

To describe the states of water in hydrogels, various models have been proposed. While the classification remains highly dependent on the involved polymers and the specific characterization method employed, there is an increasing body of research – particularly for hydrogels functionalized with hydroxyl (–OH) groups – that supports a three-state model [13]. This approach distinguishes between strongly bound water that is tightly associated with polymer chains, an intermediate weakly bound water that exchanges slowly with the surroundings, and free water, which is mobile and can flow through the interconnected porous network. An idealized scheme of such a structure is shown in Fig. 1. A similar structure with the three distinct states of water has been introduced before by Li et al. [18]. These states differ in mobility and in their contribution to the mechanical and transport phenomena of the hydrogel [19–21]. Furthermore, the amount of free-flowing fluid is directly related to the pore structure of the hydrogel. The following ranges can be associated with characteristic pore scales [20]: *<* 1.4 nm (primarily strongly bound), 1.4–3 nm (strongly and weakly bound), 3–50 nm (mixed, including free water), and *>* 50 nm (predominantly free water).

**Figure 1:**
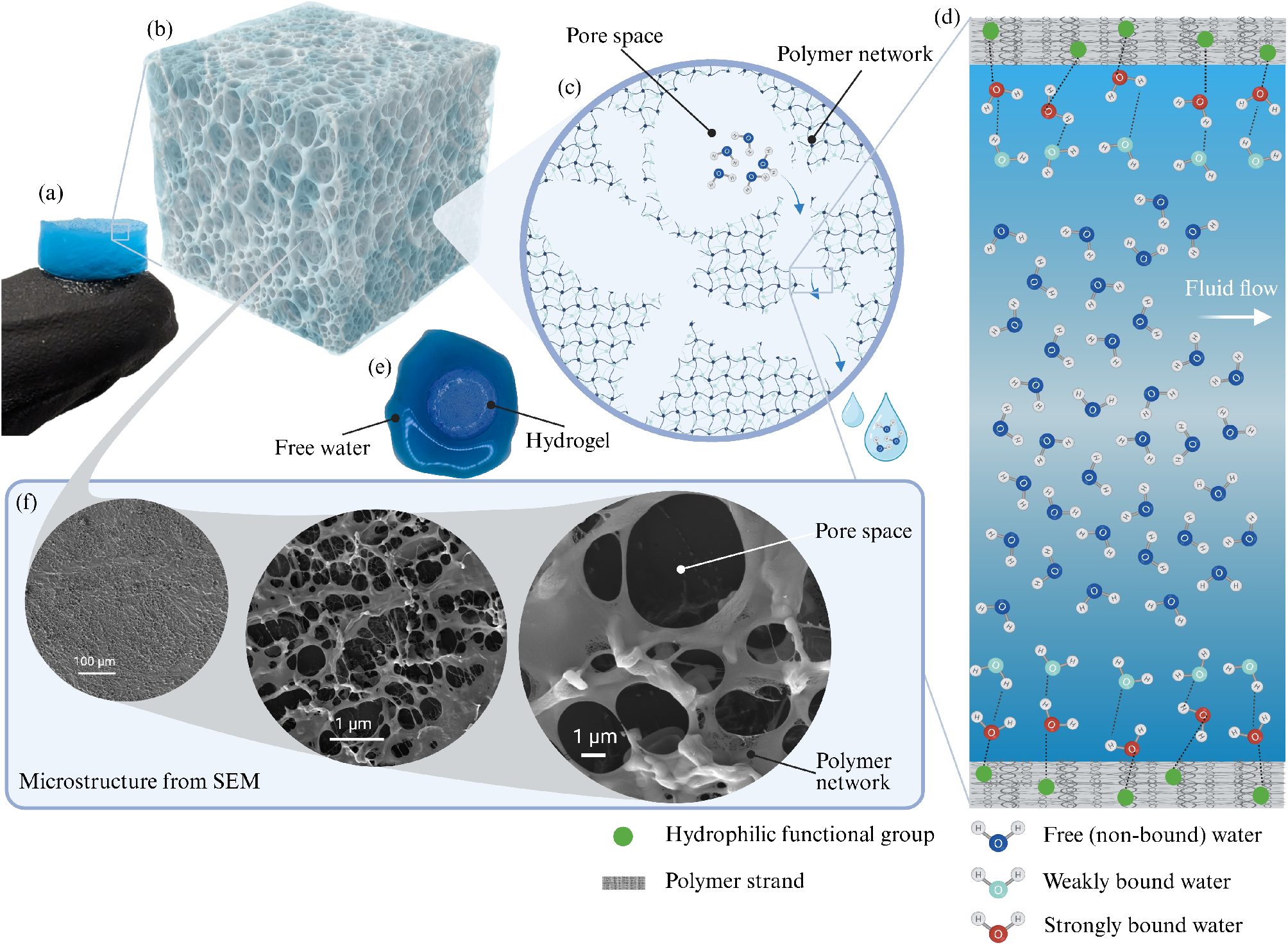
Schematic structure of a physically cross-linked hydrogel across scales: (a) macroscopic hydrogel sample with a diameter of 10 mm; (b) idealized cubic volume element showing the internal porous structure; (c) enlarged view showing the idealized structure of the polymer network (mesh) and the pore space, which contains free water; (d) close-up showing the porous structure at the molecular level, based on the ‘three-state’ water model. Adapted with permission from Li et al. [18]; (e) hydrogel sample from which free water has escaped after deformation. The box in (f) shows an actual microstructural image of the hydrogel composite recorded with cryo-scanning electron microscopy. Created in BioRender.com.

Mechanically mobile water – the fraction of free water that is displaced during deformation – plays a fundamental role in constitutive models based on biphasic or multiphasic theories [see, e.g., the monographs 22–26]. Similar to hydrated biological tissues [27, 28], hydrogels present very low stiffness and can be easily compressed. In response to an applied load, they behave as soft polymers, with nonlinear and time-dependent effects [11, 29]: at high strain rates, their mechanical behavior is dominated by rubber elasticity, whereas at longer times they can undergo large volumetric changes by exuding or absorbing water. The role of free water in the mechanics of multiphasic media lies in that it provides the material with time-dependent characteristics, governed by the chemical and physical properties of both fluid and solid phases [30]. Furthermore, the highly deformable and micron-sized porous structure of hydrogels implies that hydraulic permeability is also low, resulting in large frictional drag forces exerted by the fluid as it flows through the matrix. Hence, solid deformation and fluid flow are coupled in the mechanics of hydrogels. From this perspective, parameters such as the fraction of free water that can contribute mechanically, as well as hydraulic permeability, emerge as fundamental characteristics that require experimental identification [31, 32].

Hydrogels based on polyvinyl alcohol (PVA) are among the most used materials, particularly for engineered composite hydrogels [14, 33–35]. A common preparation method that specifically generates a macroporous network is phase templation, which is achieved, e.g., by repeating freeze–thaw cycling [36, 37]. This process produces a high fraction of free water that can be mechanically mobilized, making PVA-based, freeze-thawed composites especially suitable model systems to study how the free-water fraction depends on composition and mechanical loading [20]. Their tunable polymer concentration, simple processing, and extensive prior use as tissue phantoms further facilitate comparing different characterization methods.

Numerous experimental techniques have been introduced to investigate the porosity and free-water fraction of hydrogels. These include gravimetric methods [38], nuclear magnetic resonance (NMR) [39], and imaging methods, which are often considered a standard procedures for determining the pore characteristics of porous structures [40–43]. Among the latter, we can mention X-ray micro-computed tomography (µCT) [44] and confocal laser scanning microscopy [45, 46]. In addition, differential scanning calorimetry (DSC) has also been applied to partition non-freezing (bound) water from freezing, free water [14, 47, 48]. However, 3D image-based approaches for hydrogels and soft biological tissue typically require contrast agents or fluorescent labels such as fluorescein isothiocyanate (FITC) [49, 50], are prone to preparation artifacts, and cannot fully represent bulk volumes because of voxel-resolution limits [51]. Moreover, they assess only the static pore geometry and do not directly quantify the fraction of fluid that can be mobilized under mechanical loading, which is the quantity that enters as volume fraction in the governing equations of porous media.

On the other hand, several studies have probed the biphasic response of soft hydrogels under loading, using uniaxial compression [32, 52–58] or indentation [59–61]. In particular, Kluge et al. [53] performed drained confined compression and creep experiments on silk hydrogels, and calibrated the parameters of a linear poroelastic model, reporting permeability and fluid fraction. However, they did not directly measure total, free, or bound water fractions. Cornet et al. [57] performed confined compression in soy–protein gels and meat analogues, quantifying the expelled fluid from volume change and analyzing release kinetics. However, the free-water fraction was not separated from the total water content. Notably, both studies used protocols involving multi-step loading.

To close the current gap, we introduce a mechanical protocol based on stepwise, bottom-drained confined compression-consolidation to quantify the mechanically releasable free-water fraction. This protocol provides a direct, mechanics-based estimate of the effective porosity that is relevant for biphasic models and applications. Within the framework of the introduced three-state model of water in hydrogels, this refers to the unbound, mobile fraction that is capable of flowing through the interconnected porous network. The stepwise protocol additionally enables the identification of states of equilibrium for the material, where the response is governed by the properties of the solid matrix. In addition, we directly compare the free-water fraction derived from the incremental consolidation to two complementary metrics of porosity, namely: (i) total water content obtained from dehydration and gravimetric analysis, and (ii) surface porosity derived from cryo-scanning electron microscopy (SEM) image analysis. As a model system, we use composite, freeze-thawed PVA-based hydrogels that have proven suitable as brain phantoms [1, 62, 63], calibration materials [61], and for soft-tissue biomimicking interfaces [64]. The composite material can mechanically mimic soft tissue over large regions [65], is compatible with a variety of cells, and has been shown to be 3D-printable [66]. The chosen phase-templating freeze–thaw protocol produces an interconnected, micron-sized pore architecture (Fig. 1), whose clear morphological distinction provided the motivation for the present work. We study three formulations with increasing polymer concentration to span a practical range of stiffness and pore connectivity, and we directly compare the porosity metrics and the mechanical response across compositions.

The paper is structured as follows. We provide a compact background on aspects of the continuum theory of biphasic media in Sec. 2, which is needed to motivate the development and interpretation of the experimental methods. Experimentally, we introduce a custom bottom-drained uniaxial consolidation rig that reproduces a well-defined boundary value problem (BVP) of the biphasic theory in Sec. 3.2. Two mechanical protocols are applied: a stepwise stress–stretch protocol to establish steady-state points, and an incremental consolidation protocol to quantify displaced fluid volumes up to compaction. Section 3 is completed by methodological details about the gravimetric analysis and cryo-SEM image analysis and processing. We present and discuss the equilibrium behavior of the material in Sec. 4.2 and derive the free-water fraction at compaction in Sec. 4.3. Results from gravimetric analysis providing the total water content (Sec. 4.1), analysis of cryo-fixated cross sections (Sec. 4.5-4.6), and image-derived surface porosity (Sec. 4.4) are presented, and the three metrics compared, discussing discrepancies in terms of connectivity, closed versus open pore space, and sub-resolution polymer mesh contributions. Finally, implications for model calibration and mechano-mimetic hydrogel design are outlined in the Conclusion (Sec. 5).

## 2 Continuum Theoretical Background

This section outlines some fundamental elements of the classical continuum theory of soft porous materials. It focuses on the assumptions that led to the identification of the soft hydrogel as a biphasic medium, as well as the considerations that guided the development of the experimental protocols described in this article.

### 2.1 Hydrogels as biphasic media

A hydrogel in its swollen state can be viewed as a mixture of cross-linked polymer chains and water. In the theory of mixtures, the individual phases are treated as overlapping continua that can exchange mass, momentum, and energy with each other, while the mixture as a whole behaves as a pure substance. The fundamental balance laws are formulated for each phase and then summed to derive the corresponding principles of the mixture, subject to specific constraints. A short summary of the fundamental equations is provided in A, and the interested reader is directed to the classical monographs by Truesdell [22] and Bowen [24], which include also a discussion on the non-trivial entropy inequality in mixtures. We want to remark that this approach is substantially different from monophasic models, such as those employed in the classical theories of poroelasticity [see, e.g., 26, 67] and swelling of gels [see, e.g., 68, 69]. However, equivalence was demonstrated in the context of a biphasic mixture and infinitesimal strains [70, 71]. Further insights into the application of monophasic and mixture theories to hydrogels can be found in the literature [72, 73].

In this work, we consider a homogeneous mixture formed by a solid phase (hereafter denoted by the superscript s) and a fluid phase (identified by f), which are assumed to be immiscible and individually incompressible [70]. For each phase *α* ∈ {s, f} we can then define the *volume fraction ϕ*^*α*^ as the local ratio of the current volume element of the *α*-th phase with respect to the current volume element of the mixture, i.e.

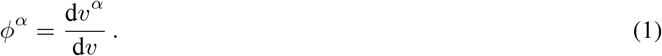

As discussed in the Introduction, the fluid phase in hydrogels is water, which can exist in different forms. In the biphasic mixture considered in this work, the solid phase defines a control domain through which the fluid phase flows, exchanging mass and momentum. Accordingly, in the mechanical response, it is essential to account only for the volume fraction of the free, non-bound water that can move through the interconnected pores of the solid network, whereas the bound water is regarded as part of the solid matrix. Under the assumption that the mixture is always *saturated*, the volume fraction of the fluid can thus be identified with the *effective porosity ϕ*^f,eff^.

Once this equivalence is theoretically established, the focus shifts to its experimental determination. In this work, we propose a method for estimating the effective porosity based on mechanical consolidation and compaction, which is described in detail in the following section. On the other hand, microstructural imaging can provide information about the fraction of pore space and its arrangement. This point acquires particular significance in the context of developing a continuum representation of the biphasic mixture. Although the classical theory of mixtures is phenomenological, meaning that the continuous macroscopic fields are not explicitly derived from averaging procedures at the microscale [74], any experimental measurement can be regarded as an averaging operation over a finite region defined by the resolution of the measurement technique and the dimensions of the sample. Therefore, a correspondence must be established between the experimentally derived quantity and the associated continuous field [23].

To formalize this concept, we introduce an averaging operation at the mesoscopic scale (i.e., the intermediate level between fluid molecules and the macroscopic scale of the porous medium) and define the *areal porosity* in the reference configuration of the solid as [75]

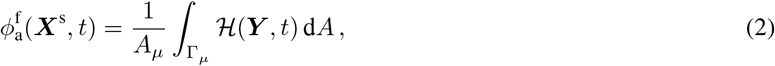

where *A*_*µ*_ denotes the area of a control domain 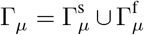 attached to a material point of the mixture, identified by its reference position ***X***^s^, and ***Y*** ∈ Γ_*µ*_ is an arbitrary point within this domain (Fig. 2(a)). The scalar function ℋ represents the fluid distribution within the control domain, defined as

**Figure 2:**
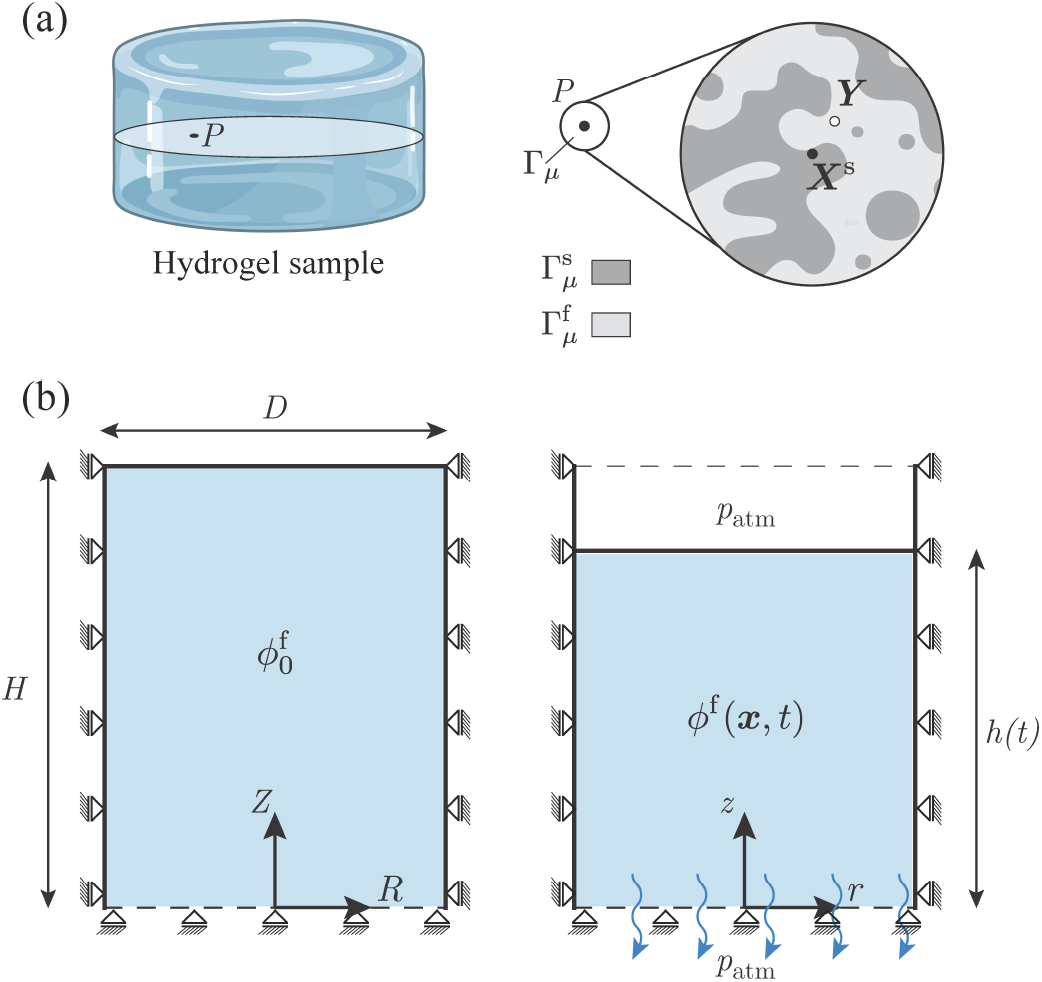
(a) Macroscale and mesoscale in the reference configuration of a hydrogel sample. At point *P* an averaging operation is defined to obtain the areal porosity field 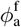 ; (b) sketch of the boundary-value problem (BVP) of drained confined compression and consolidation of a cylindrical homogeneous sample, showing the reference and current configurations. Created in BioRender.com.

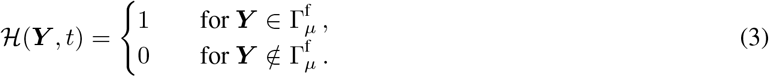

Applying this method requires an analysis of the mesoscopic structure of the undeformed hydrogel samples, which in this work is carried out following the procedures described in Sec. 3.4. According to Delesse’s law, the areal fraction of a random porous network coincides with the volumetric fraction [23].

### 2.2 Uniaxial deformation and flow

Solid deformation and fluid flow are generally coupled in the mechanics of biphasic porous media. In addition to the classical conservation laws, constitutive relationships for the individual phases and their interaction must be formulated [76]. Therefore, a long-term goal of our research is to establish simple BVPs that can be translated into experimental setups, enabling the solution of the inverse problem of calibrating the constitutive laws. To this end, we select a class of BVPs involving uniaxial deformation and fluid flow in a cylindrical domain.

We consider the swollen state of the hydrogel without deformation as the reference configuration and assume isotropy and homogeneity (Fig. 2(b)). Accordingly, we describe the mechanical response about this state and do not explicitly account for the osmotic swelling process by which the reference state is attained. To describe the solid deformation and the fluid flow in the biphasic medium, we introduce the following primary fields, expressed in cylindrical coordinates (*r, θ, z*):

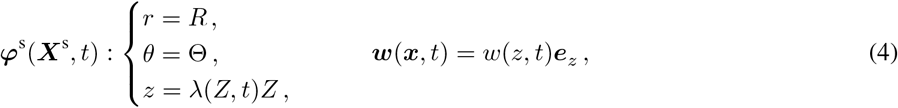

where ***φ***^s^ denotes the motion of a material point ***X***^s^ of the solid phase, ***x*** is the current position of a point in the mixture, and 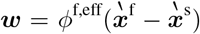 is the *discharge velocity* vector, defined from the velocities 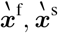 of the fluid and solid phases, respectively. This term represents the volumetric flux of fluid relative to the solid, per unit area of the mixture and per unit time. The longitudinal stretch *λ* = *λ*(*Z, t*) is used in this work to quantify the uniaxial deformation of the solid, which is in general inhomogeneous and time-dependent, as highlighted by its dependence on the reference longitudinal coordinate of the point *Z* and the time *t*. Notably, Eq. (4)_1_ shows that the deformation gradient depends only on the change in porosity [77], since

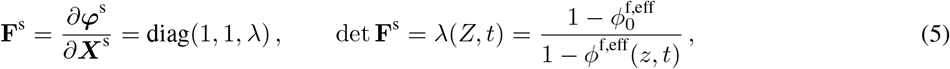

where 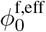 is the effective porosity in the reference configuration. In a homogeneous mixture, this is assumed spatially uniform; however, due to the inhomogeneity of the deformation, the current effective porosity *ϕ* ^f,eff^ can be non-uniform with respect to the longitudinal coordinate.

Here we focus on the case in which the upper boundary of the cylindrical sample is compressed by a rigid, impermeable plate under a prescribed displacement, while the lateral boundary is frictionless and restrains displacement and fluid flow, and the bottom boundary is rigid and drained (Fig. 2(b)). In the literature, this is known as confined compression experiment with free draining, in short drained confined compression. As the plate moves downward, the solid deforms, driving fluid out of the material through a reduction in porosity. If the displacement is held fixed, *h*(*t*) = *h*_0_, a *consolidation* process occurs in which the total reaction force relaxes over time until equilibrium is reached asymptotically, i.e. for *t*→ ∞. In practical terms, this conditions is approximated after a sufficiently long holding time *t*^eq^. The equilibrium, steady state is characterized by zero fluid flow and homogeneous deformation throughout the sample, i.e., *w*(*z, t*^eq^) = *w*^eq^ = 0 and *λ*(*Z, t*^eq^) = *λ*^eq^ = *h*(*t*^eq^)*/H* (see A.3). At this stage, the reaction force on the upper plate corresponds to a homogeneous longitudinal stress in the solid skeleton, with the fluid completely stress-free. In contrast, the transient is marked by highly inhomogeneous fields, and the reaction force includes evolving solid and fluid contributions that depend on the constitutive behavior of each phase.

It is important to note that the homogeneous state is achieved asymptotically, analogously to the equilibrium state of a viscoelastic solid. Given the practical challenges in achieving this state under continuous loading, it appears reasonable to define a multi-step protocol, in which each loading step *i* = 1, …, *N* corresponds to a consolidated configuration of the sample with *λ*^eq,*i*^ = *h*(*t*^eq,*i*^)*/H*. Furthermore, this protocol allows the *point of compaction* to be reached, defined as the state at which the total free water has been expelled and the incompressibility of the solid phase prevents any further deformation [78]. Assuming phase incompressibility, the stretch *λ*^eq,*N*^ at this point, which quantifies the total volume change in the biphasic material, yields the initial effective porosity according to

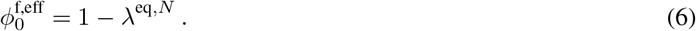

## 3 Experimental Tests

### 3.1 Preparation of composite hydrogels

Composite hydrogels were prepared following protocols adapted from previous studies [1, 64]. Polyvinyl alcohol (PVA) (MW 146 000-186 000, 99+ % hydrolyzed) and Phytagel (PHY) granulates were obtained from Sigma-Aldrich and used without further modification. PVA and PHY were dissolved in deionized water (DIW) under continuous stirring and heating until fully solubilized. Three composite hydrogel (CH) formulations were produced with varying polymer concentrations (w/w%): CH1 (2 % PVA, 0.3 % PHY), CH2 (3 % PVA, 0.4 % PHY), and CH3 (4% PVA, 0.6 % PHY). These specific ratios were selected based on preliminary mechanical characterization to span a relevant stiffness range representative of soft biological tissues [64]. Following dissolution, the solutions were cooled to room temperature and subjected to a single freeze–thaw cycle to achieve physical cross-linking of the double network structure [1]: freezing at −20 ^*°*^C for 24 h, followed by thawing for 12 h under standard laboratory conditions. No additional cross-linking agents were used. The resulting hydrogels were stored hydrated in DIW with 0.03 vol% triarylmethane dye, a standard food coloring. Swelling of the sample due to the addition of the dye or any particles could not be detected. For material characterization, hydrogel discs were extracted from bulk material, trimmed to a height of *H* = 5 ±1 mm, and then cut with a commercial biopsy punch to a nominal diameter of *D* = 10 mm.

### 3.2 Confined compression tests

#### 3.2.1 Experimental setup

The mechanical tests were conducted using a triaxial testing device (ZwickRoell Testing Systems GmbH, Fürstenfeld, Austria) previously described by Sommer et al. [79]. The setup was extended with a custom-built test rig based on the fundamental principles of an oedometer. The rig enables uniaxial compression and consolidation of cylindrical samples inside a transparent, rigid chamber (Fig. 3a). The upper piston applies a vertical load, while the bottom end features a permeable support structure composed of a rigid grid holder, a stainless steel mesh, and a lint-free cellulose filter. This assembly permits the controlled expulsion of fluid during compression. The expelled fluid is absorbed by a rayon-cotton compound beneath the chamber, ensuring minimal reabsorption. Filter retention and fluid permeability were selected to exceed the expected fluid flow rate from the sample, preventing congestion effects as reported in similar tests [80]. A filtration rate of approximately 10 mL/min was ensured, significantly above the rates reported for this material [61]. A small clearance between piston and cylinder is required to avoid measuring parasitic friction forces. Here, the fixture was designed with a gap of approximately 0.1 mm, maintaining a lubricating water film between the surfaces. This configuration minimizes shear at the walls and is standard practice in confined compression setups [53–55, 57]. The setup allows time-resolved measurement of force and displacement, enabling quantification of the volumetric changes due to expelled fluid. Since the absolute displaced fluid volume is small (less than 1 mL over several hours in trial tests), and thus prone to evaporation, the longitudinal displacement has been used to derive the change in fluid content indirectly. This method assumes incompressibility of the phases involved, an assumption introduced in Sec. 2.2 and further discussed in Sec. 4.1.

**Figure 3:**
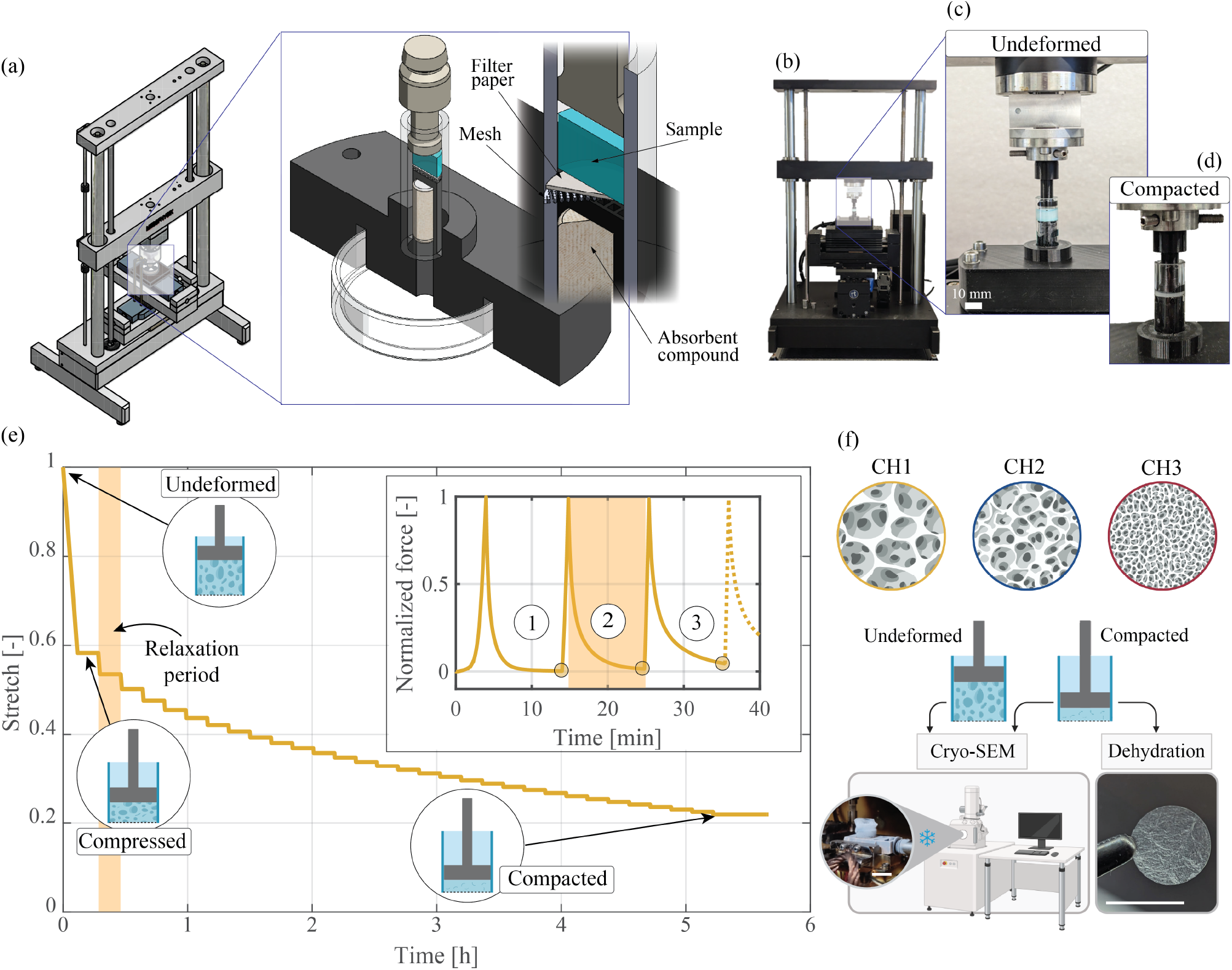
Overview of the methods employed for the experimental assessment of porosity in the hydrogel composites: (a) sketch of the confined compression testing device, with an enlargement of the custom-built test rig highlighting details such as the sample (blue), the filter paper, the metal mesh, and the absorbent compound to collect expelled fluid; (b) image of the setup; (c) saturated sample in the undeformed state; (d) compacted sample; (e) loading protocol of the incremental consolidation experiment in the form of macroscopic stretch-time curve. Highlighted are three main phases – the *undeformed* state at *t* = 0, the first *compressed* state, which results as soon as a threshold force value is reached, and the *compacted* state after a series of consolidation steps with an extended holding/relaxation time. The shaded area highlights the relaxation period in the main figure and in the inset, which shows the force response in a normalized form. The first three consolidation steps, indicated by (1), (2) and (3) are shown; (f) cryo-scanning electron microscopy imaging and dehydration workflow for the three composite hydrogels (CH) used in this work. Created in BioRender.com.

#### 3.2.2 Testing protocols

The samples were weighed three times before loading in the consolidation cell. The sample height was determined by lowering the upper piston at a rate of 2 mm/min until a contact force of 2 mN was reached, taking into account possible deformations due to the sample’s own weight, particularly with the softer hydrogel composition. This sample geometry in the confined chamber was then taken as a reference for subsequent analysis. All test protocols were displacement-controlled with a data acquisition rate of 1 Hz. In the loading phase, a constant speed *v* of the piston is defined and adjusted based on the initial sample height *H*, with 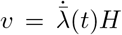, where 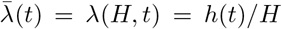 is defined as the macroscopic stretch. Macroscopic stretch rates of 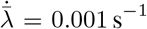 for CH1 and CH2 and 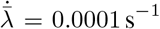 for CH3 were used (Fig. 2(b)). The reaction force recorded by the load cell in the longitudinal direction is normalized by the cross-sectional area of the sample, *A* = *πD*^2^*/*4, providing the nominal stress at the top plate in the longitudinal direction. Representative images of the experimental setup can be seen in Fig. 3(b)-(d).

We adopted two distinct protocols. The first, referred to as *stepwise compression*, consisted of subsequent steps of compression-relaxation at predefined stretch ratios 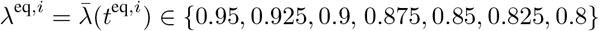 for CH1 and CH2, and *λ*^eq,*i*^ ∈ {0.95, 0.925, 0.9, 0.875, 0.85} for CH3. The step at 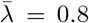 was excluded for CH3 because of excessive reaction force, which could damage the load cell. The relaxation times were adjusted such that an equilibrium state could be achieved. For CH1 and CH2, the seven steps were performed with a constant relaxation time of Δ*t*^*i*^ = 600 s. For CH3, the relaxation times were adjusted to Δ*t*^1^ = Δ*t*^2^ = 900 s, and Δ*t*^*i*^ = 1500 s for the remaining steps. The longitudinal reaction force was normalized by the initial cross-sectional area of each sample, providing the nominal stress of the solid at each equilibrium point. The loading protocols and corresponding responses are presented in Sec. 4.2.

The second protocol, referred to as *incremental consolidation*, was designed to quantify the total amount of free water in the hydrogels, i.e., the maximum volume of fluid that can be displaced through consolidation. In this protocol, the load was gradually increased and the sample compressed until a threshold force of 2 N was reached, followed by a holding period of 600 s. A representative stretch history is presented in Fig. 3(e). This incremental loading-holding sequence was repeated until no further compression was possible. This indicated that a compaction point had been reached while maintaining a moderate nominal stress of 25 kPa. The macroscopic stretch at this point was used to estimate the free water volume based on the assumption of incompressibility of the solid matrix. Across all hydrogel compositions, a maximum of 31 steps was found to be sufficient to reach compaction. Results of the incremental consolidation experiment are shown in Sec. 4.3.

For the *stepwise compression, n* = 6 samples of CH1, *n* = 9 samples of CH2, and *n* = 5 samples of CH3 were analyzed. For the *incremental consolidation* protocol, *n* = 8 samples of CH1, *n* = 9 samples of CH2, and *n* = 10 samples of CH3 were analyzed. After mechanical testing, samples were prepared for dehydration. Unless otherwise stated, all mechanical tests were performed at a temperature of 22 ± 2 ^*°*^C.

### 3.3 Dehydration and total water content

To assess the total amount of water in the hydrogels in line with the three-state model of water [13], hydrogel discs were fully dehydrated following the completion of mechanical testing. The procedure followed the gravimetric method described in earlier work [61]. In brief, samples were dehydrated over 120 h at 22 ±2 ^*°*^C. The mass was measured in triplicate both before and after dehydration, allowing the derivation of the amount fluid under the approximation of similar bulk densities for solid and fluid phases, i.e. *ρ*^f^ ≈*ρ*^s^, and *ρ*^s^ = 1.0 g/cm^3^ in the fully fluid-saturated state [81]. A total of *n* = 11 discs of each hydrogel composition were dehydrated. No distinction was made with respect to the mechanical protocol applied to the hydrogel sample before dehydration.

### 3.4 Cryo-scanning electron microscopy

The microstructure of the composite hydrogels was examined using cryo-SEM. Cylindrical samples were rapidly frozen in slush liquid nitrogen and transferred into a cryo-preparation chamber (Quorum PP3010T, Quorum Technologies, UK), which was directly coupled to a GEMINI Sigma 500 SEM (Carl Zeiss, Germany) equipped with a nitrogen-cooled stage. To avoid influences on the subsequent analysis due to contamination on the sample surface, each sample was fractured under vacuum in the cryo-chamber followed by controlled sublimation to reveal the bulk hydrogel structure. Subsequently, samples were sputter-coated with a thin layer of palladium and transferred under vacuum into the SEM imaging chamber. Imaging was performed using a 2 kV and 5 kV acceleration voltage with a secondary electron detector. Cryo-SEM was conducted for both undeformed and consolidated/compacted hydrogel samples. Post-consolidation, samples were mechanically fixed in their final loading configuration, rapidly frozen and stored at −80 ^*°*^C to preserve their structural state. Samples were continuously maintained at cryogenic temperatures until imaging. All samples were prepared and imaged following the same standardized cryo-SEM protocol described above. The processes describing imaging of undeformed and compacted samples of the three hydrogel compositions and the subsequent dehydration are summarized in Fig. 3(f).

#### 3.4.1 Image processing

The SEM images were processed and segmented to analyze the surface porosity and to investigate its relationship to the observation scale. Surface porosity was defined as the ratio of pore area in reference to a region of interest (ROI). Here, the ROI refers to the area of the original image, or a section of it. The open-source software *ImageJ* with the toolbox *Analyse Particles* was used to derive the surface porosity [82] as a reference. A detailed description of such an analysis is presented in [42] and in the Supplementary Material. In brief, original images were scaled according to magnification to preserve structural dimensions. A grayscale threshold was applied to binarize the images, segmenting the dark background (frozen bulk water) from the brighter polymer network exposed through controlled sublimation. The surface porosity was quantified as the ratio of pore area to total area within a defined ROI, using the ImageJ parameter *%Area*. For the quantitative analysis, *n* = 5 images with varying magnification per hydrogel composition were processed. The images are summarized in the Supplementary Materials.

Porous hydrogel do not necessarily have a homogeneous internal structure, and the surface porosity can therefore vary locally [83]. In addition to evaluating the overview images, we analyzed the potential influence of the image area on the surface porosity using a Matlab-based algorithm. Starting from the original rectangular image, we chose the largest square window, of size *L*× *L*, that fits inside the image and raster-scanned that square in row-major order (left →right, top →bottom), advancing one pixel at a time and only evaluating positions where the full square lies inside the image. For every window position, the algorithm computes the local porosity as the fraction of pore pixels within the square window. The window size is then reduced by one pixel in both dimensions and the entire raster scan is repeated; this process continues stepwise until the window size is one pixel. The data for all possible sizes and positions were then averaged and presented in the form of median and inter-quartile range (IQR). The window size is mapped to a physical structure size via calibration of the imaging data. With *p* representing the pixel density of an individual image expressed in pixels per µm, the structure size *S*_*L*_ is

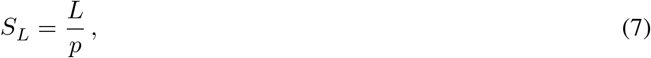

and the corresponding normalized window area fraction *A*_*L*_ is

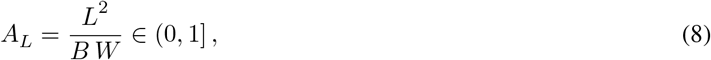

where *L* is the length of the current window, and *B* and *W* are the dimensions of the original image. Plotting the median and IQR versus structure size and versus normalized area thus reveals how surface porosity evolves with the observation scale.

### 3.5 Statistical analysis

Pairwise comparisons of the medians between the two methods (surface porosity vs. effective porosity) were performed within each composition (CH1, CH2, CH3), and, where relevant, across compositions (CH1 vs. CH2, CH1 vs. CH3, CH2 vs. CH3). Statistical significance was set at *p <* 0.05. To quantify the magnitude of between-method differences, Cliff’s delta *δ* was calculated as a nonparametric measure of effect size for median differences. We additionally reported the relative difference between methods in percent, defined as the absolute difference between their median values divided by the effective porosity.

## 4 Results and Discussion

### 4.1 Gravimetric analysis: total fluid fraction and the assumption of incompressibility

The gravimetric analysis provided an estimate of the total fluid fraction in the three hydrogel compositions. Measurements of the mass of each sample before compression and after compression–dehydration were used to determine the total solid fraction *ϕ*^s,tot^, as shown in Fig. 4. Assuming saturation, the corresponding total fluid fractions *ϕ*^f,tot^ = 1 −*ϕ*^s,tot^ were found to be 0.976 for CH1, 0.965 for CH2, and 0.955 for CH3. The obtained total fluid fractions lie within a remarkably narrow range, confirming that all hydrogels have a very high water content. These values are in the upper range of those reported for other PVA-based hydrogels [14, 84] and confirm that all three formulations are highly water-dominated composites. This observation highlights the exceptional water-binding capacity of these materials, which is an important property for their applications [13].

**Figure 4:**
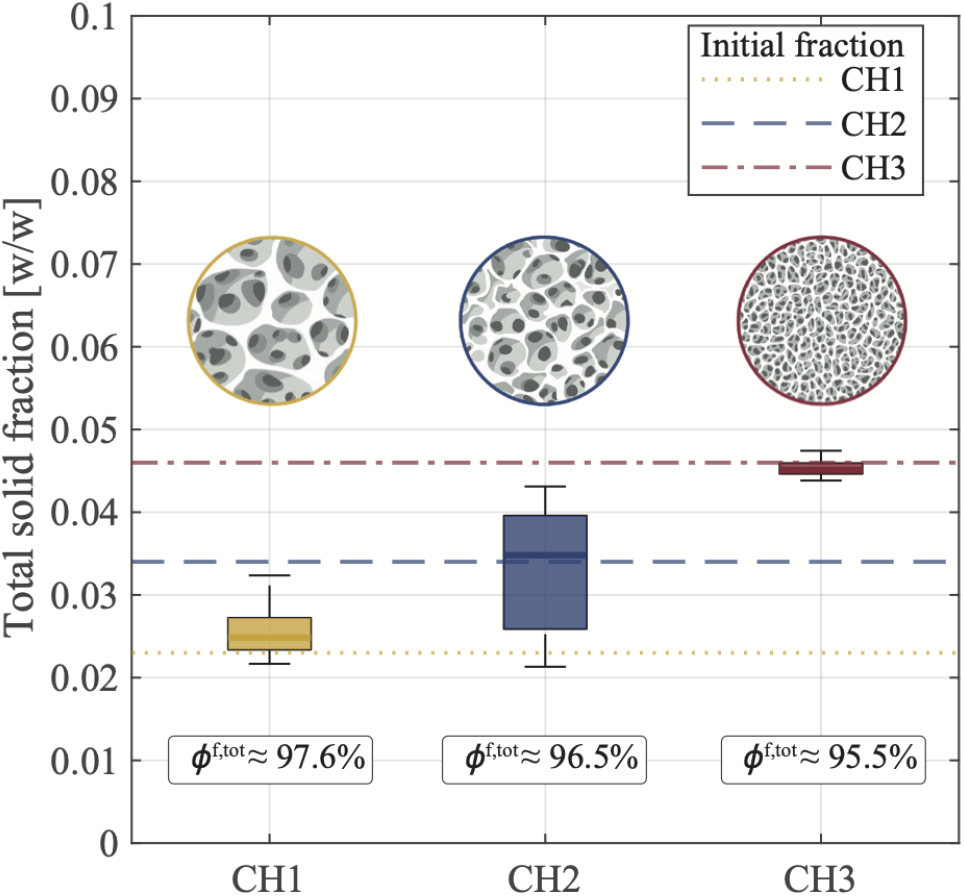
Total solid volume fraction for the three hydrogel compositions obtained from the gravimetric analysis on dehydrated hydrogel samples. Boxes depict the median values with *n* = 11 for each composition. Dashed lines indicate the nominal values, which are based on the polymer fraction defined in the preparation protocols. The total fluid volume fraction *ϕ*^f,tot^ displayed is computed from the median assuming saturation. Created in BioRender.com.

From a mechanical perspective, the high water content of hydrogels is often cited to justify the assumption of material incompressibility [85, 86]. Several authors [see, e.g., 87–89] have attempted to support this assumption by providing experimental measurements of the Poisson’s ratio, although these analyses usually do not account for the large strain regime. It is important to clarify, however, that incompressibility strictly applies to the undrained response of the material, whereas large volume changes can occur as a result of fluid drainage [69]. In contrast, incompressibility of individual phases, both fluid and solid, is a fundamental assumption in our methods, which allowed us to employ uniaxial consolidation to derive the total amount of free-flowing fluid. As clarified in Sec. 3.2, direct quantification of the expelled fluid was avoided to minimize potential errors due to evaporation during the long testing protocols. Instead, the displaced fluid volume was derived from the total volume change under confined compression, see Eq. (6), which requires the assumption that both solid and fluid phases are incompressible. While water can be safely treated as incompressible, this assumption is less trivial for the polymeric solid phase. In the absence of reliable experimental data, we consider this a reasonable approximation, since any compressibility of the solid network is expected to be negligible compared to the large volume changes induced by fluid outflow during consolidation. This assumption is also widely adopted in classical theoretical studies on hydrogels [see, e.g., 69, 90–92].

### 4.2 Stepwise compression: relaxation and equilibrium points

The results of the stepwise, displacement-controlled compression are presented in Fig. 5, showing the loading protocol together with the force response for the three hydrogel compositions. Each loading step leads to a pronounced peak in the reaction force, followed by a gradual relaxation during the holding phases. The nominal stress is determined by normalizing the reaction force by the cross-sectional area of the samples. It should be emphasized that the reported stretch and nominal stress are representative of a material point in the immediate vicinity of the cover plate, as the deformation state – apart from the steady-state points – is inhomogeneous. Two limiting responses can occur in this loading scenario: (i) no fluid mobilization, in which the reaction force remains constant during the holding phase or exhibits only a relaxation associated with the intrinsic viscoelasticity of the polymer matrix; (ii) fluid drainage, producing a large volumetric change and a pronounced peak–relaxation behavior upon compression. The results presented in Fig. 5, which demonstrate a remarkable force relaxation, strongly suggest that the mobilization and drainage of free water from the bulk composite is the dominant mechanism.

**Figure 5:**
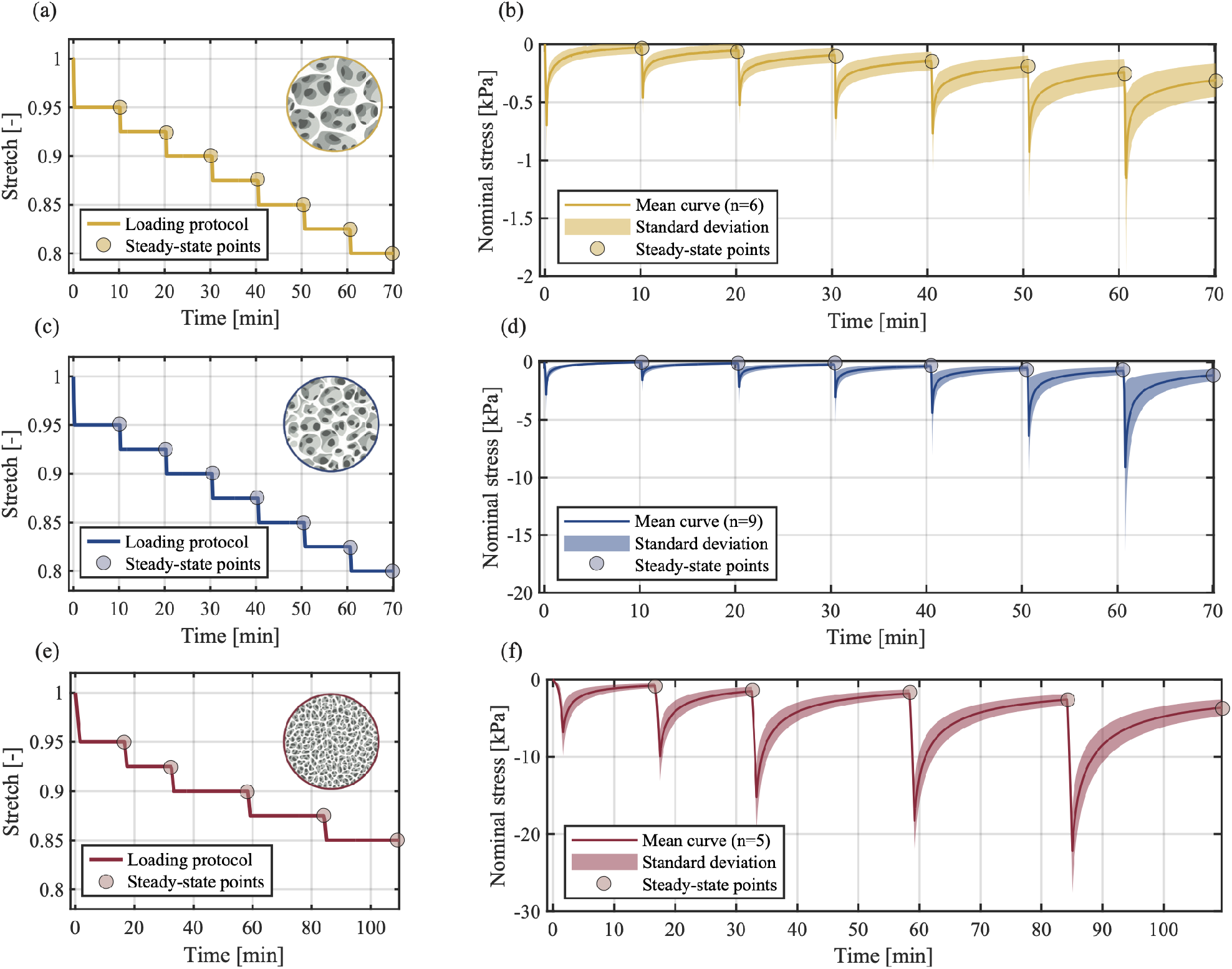
Stepwise compression: (a),(c),(e) loading protocols for the three hydrogel compositions (CH1–CH3), showing the macroscopic stretch history 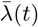 and the steady-state points at *λ*^eq,*i*^; (b),(d),(e) corresponding mechanical responses in terms of normalized longitudinal reaction force recorded during each protocol. Created in BioRender.com.

Both peak and equilibrium forces scale notably with polymer content (CH3*>*CH2*>*CH1), i.e., compositions with higher polymer content show higher peak and steady-state forces. This becomes particularly evident when the steady-state points are plotted against the corresponding macroscopic stretches (Fig. 6). At the time each steady-state point is extracted, transient and dissipative processes have subsided, and the measured normalized force represents the equilibrium stress 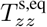 in the solid skeleton. According to the theoretical considerations developed in Sec. 2.1, these are *equilibrium points* characterized by a homogeneous stretch *λ*(*Z, t* ^eq,*I*^) = *λ*^eq,*I*^, and zero fluid flux, *w*(*Z, t*^eq,*i*^) = 0. In other terms, the equilibrium curve that can be obtained by ideally connecting these points describes the response of the porous solid and can serve as the basis for identifying the mechanical parameters of the solid skeleton. As noted in the seminal work of Treloar [29], this response is that of a compressible monophasic material. The definition of equilibrium is consistent with the long-time limit introduced in the classic swelling theories [see, e.g., 69], which corresponds to a homogeneous chemical potential in the problem domain, as determined by the solvent. Because our theory is purely mechanical and the solvent is a barotropic fluid, the chemical potential is expressed in terms of the hydrodynamic pressure *p* and the fluid density *ρ*^f^. The experiment investigates hydrogel consolidation from a swollen reference state, neglecting the role of osmotic pressure in achieving that swollen state. As explained by Chester and Anand [90], a consolidation experiment describes the drainage of fluid under mechanical compaction of the solid, starting from a swollen equilibrium state in which the fluid volume fraction is 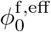, and progressing asymptotically via a progressive decrease of *ϕ*^f,eff^ until a steady state is reached. Notably, a step-wise protocol allows the assessment of multiple steady-state points and associate fluid volume fractions with solid deformation. Unlike the steady state, the transient response arises from the individual mechanical behaviors of the fluid and solid, as well as their interaction.

**Figure 6:**
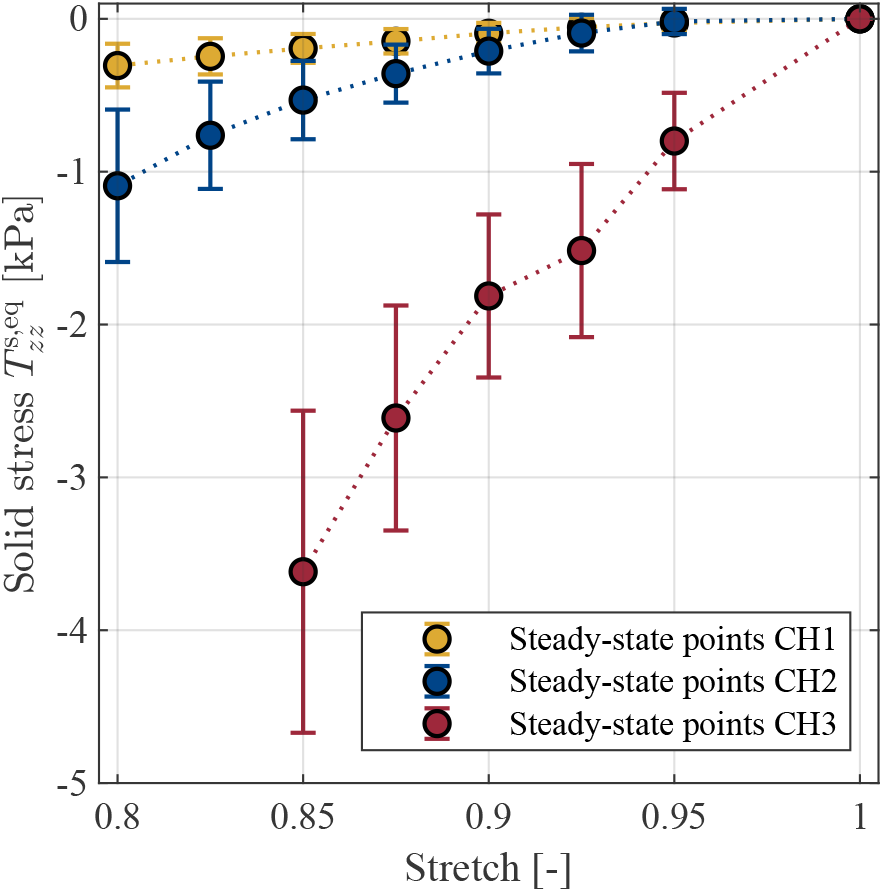
Mean equilibrium curves obtained from stepwise compression tests. The solid stress 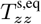 at the steady-state points is plotted against the longitudinal stretch *λ*^eq,*i*^ for the three hydrogel compositions: CH1 (yellow), CH2 (blue) and CH3 (red). Each marker represents the mean of the normalized force extracted at the end of the relaxation period; the vertical error bars show the standard deviation. For clarity, the markers are connected by dotted lines.

Calibrating a constitutive model for this coupled response requires identifying a suitable *permeability law*. Within mixture theories, this is related to the interaction force between fluid and solid phases. Rather than modeling swelling as a diffusion of solvent molecules through the gel, this term captures the frictional interaction of convective flow through the porous skeleton. Given the large volume of electro-neutral fluid in the swollen hydrogels considered here, this latter mechanism is likely dominant [73]. The key ingredient of a permeability law in soft porous materials is the reduction of permeability as porosity decreases, here identified as the effective porosity *ϕ*^f,eff^, tending towards zero at the compaction point [93]. Instead of pre-selecting a specific form, such as a permeability of the Kozeny-Carman or exponential type [27], we can integrate a very general dependence on the strain and the relative fluid-solid velocity. The calibration of such a constitutive law, requiring an additional experimental setup, is currently under development and will be addressed in a follow-up study.

The current framework does not include diffusion of reactive chemical species within the solvent, which can, e.g., be relevant to describe the mechanics of active polymer gels [94, 95]. In addition, there might be situations in which both mechanically-driven convective flow and chemically-driven diffusion coexist, and therefore both hydrodynamic and osmotic pressures are relevant [96]. The framework of mixture theories provides a rigorous basis for multi-component mixtures, where neutral or charged solutes diffuse in the solvent phase [97, 98]. In such cases, the permeability relationship would also depend on the solute concentration and the diffusivity of the solute in the solvent. Given the focus of this paper on mechanical experiments, this possibility is left as a possible future extension.

We further compare the response of the three hydrogel compositions qualitatively. In terms of relaxation behavior, CH1 and CH2 show a rapid, ‘L-shaped’ decay immediately after the peak, whereas CH3 exhibits a more gradual relaxation. In terms of poroelastic relaxation, this ‘L-shaped’ response reflects rapid fluid release in softer, more porous networks, while the slower decay in CH3 is consistent with greater flow resistance, reduced permeability, and a broader distribution of relaxation time scales. The same trend is evident in the increasing peak forces across the three compositions, indicating higher frictional resistance between solid and fluid phases during compression.

It is worth recalling that any additional viscoelastic process happening in the solid network cannot be distinguished with the present protocol. Viscous dissipation could also arise at intermediate relaxation timescales during the transient phase due to weakly bound water. Although several authors have proposed methods to separate relaxation due to fluid flow from other viscoelastic effects [see, e.g., 59, 99–101], to the best of our knowledge they relied on simplified assumptions that are not developed within the fully coupled nonlinear theory of porous media. With the protocol proposed we are able to identify equilibrium points at which both dissipative processes relax, since longer periods of time are required for fluid drainage in macroscopic hydrogel samples [102].

### 4.3 Incremental consolidation: free-water fraction and point of compaction

The free-flowing fluid in hydrogels is defined as the fraction of water that can flow through the interconnected porous space. In the simplified biphasic model adopted in this study, repeated mechanical loading and drainage progressively expel this free water, ultimately reaching a state in which virtually all free fluid has been removed, known as point of compaction. Under the assumption of phase incompressibility, the volumetric change associated with this consolidation process can be quantified, allowing estimation of the initial effective porosity 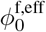 from the point of compaction, see Eq. (6).

Figure 7(a) presents the stretch–time response for the three hydrogel compositions subjected to incremental consolidation. As described in Sec. 3.2.2, vertical displacement was applied at a constant rate until a prescribed threshold force was reached, after which the piston position was held constant until all dissipative effects subsided. The cycle was then repeated until the incremental change in stretch became negligible or fell below the instrument resolution. A representative force-stretch curve is shown in Fig. 7(b), where the reaction force is normalized by the threshold force, such that the ordinate ranges between zero and one. Initially, the sample is in a saturated state at *λ*(*Z*, 0) = 1. During the first loading step, the piston is displaced at constant velocity, and the water is expelled with relatively little resistance. This leads to a gentle force-stretch slope up to approximately 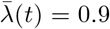. The continued compression and drainage cause a progressive collapse of the pore space, thereby decreasing the permeability and causing the compression force to rise more steeply until the threshold is reached, as indicated by the gray-shaded region in Fig. 7(b). This peak is followed by a holding period in which the force decays at constant stretch before the next loading cycle begins.

**Figure 7:**
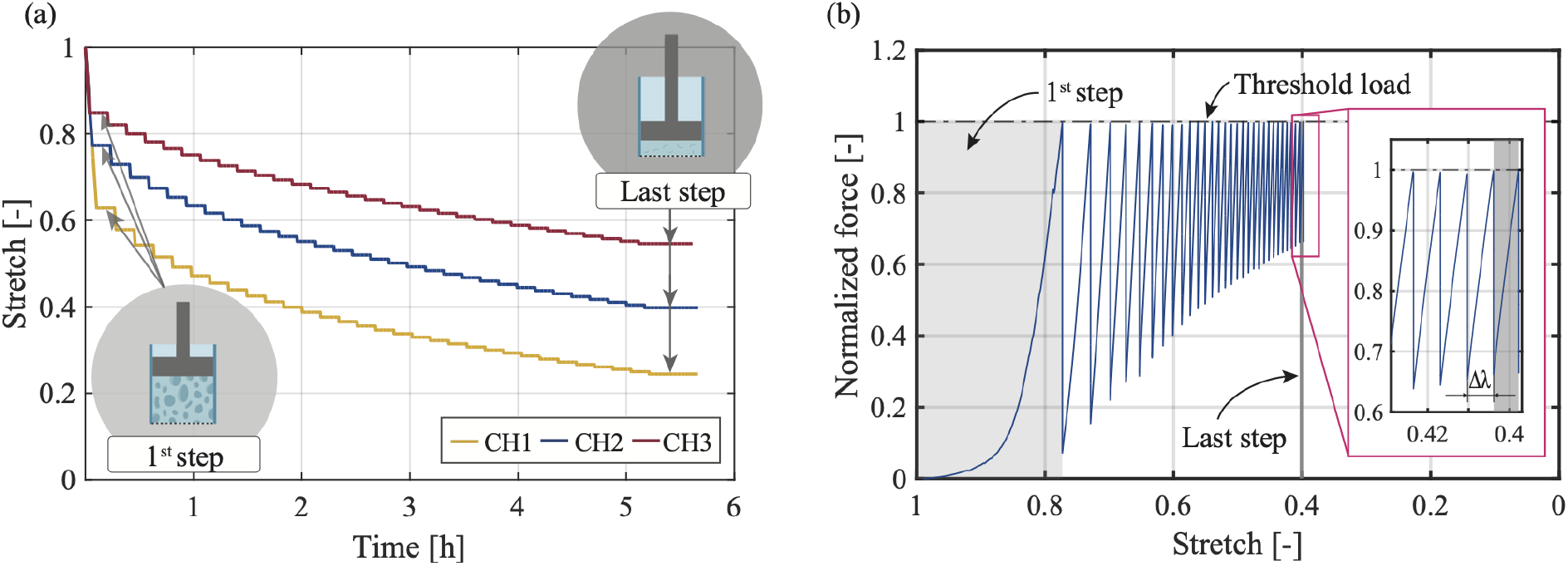
Incremental consolidation: (a) stretch-time response for the three compositions tested with identical loading protocols. Three representative datasets are shown. The varying porosity and stiffness of the hydrogels result in the depicted distinct stretch-responses; (b) Representative curve showing the normalized force over the stretch. Incrementally, the sample is compressed to the threshold load (here normalized to one) until the change at stretch 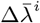 reaches the resolution limit of the equipment. The range of the first loading step and the range of the last recorded loading step are highlighted with a shaded area for comparison. Created in BioRender.com.

The curve in Fig. 7(b) captures the physical process of compaction through three concurrent trends: (i) the loading curve becomes increasingly steep, (ii) the distance between successive threshold events decreases, and (iii) the post-relaxation load increases. Trends (i) and (ii) reflect the decreasing permeability of the material, i.e., it becomes progressively harder to drive fluid through the porous network. Trend (iii) indicates the increasing load-bearing contribution of the polymer matrix, which is progressively compressed. The inset in Fig. 7(b) shows that the incremental stretch change 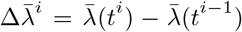 diminishes towards the end of the protocol and finally falls below the resolution of the instrument within the gray-shaded area at approximately 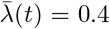 for this representative curve. This behavior indicates that the porous network has reached mechanical compaction. The corresponding stretch 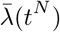 is assumed to correspond to a final steady state of the material, hence is used to determine the volume fraction of free water according to Eq. (6). The results for all three hydrogel compositions are presented in Fig. 8, which reports the volume fraction of free water together with the total fluid fraction obtained from dehydration experiments and the bound water fraction calculated as the difference. The free-water fractions were 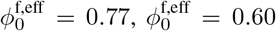, and 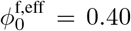, for hydrogels CH1, CH2 and CH3, respectively (median values). The sample sizes were *n* = 8 (CH1), *n* = 9 (CH2), and *n* = 10 (CH3). The amount of free-flowing fluid is in relation with the porous network of the hydrogel. Solid-phase templation via thawing, as employed in this work, produces open, macroporous architectures. In such systems, with physical pore sizes exceeding 50 nm, the free-water fraction generally dominates in the total fluid content [20], while the combined strongly and weakly bound fractions typically contribute less than 10% of the total water [12]. This behavior is evident for CH1 and CH2. In contrast, for CH3 the polymer concentration is sufficiently high to present a larger density of hydrophilic functional groups (Fig. 1), shifting the balance toward bound water.

**Figure 8:**
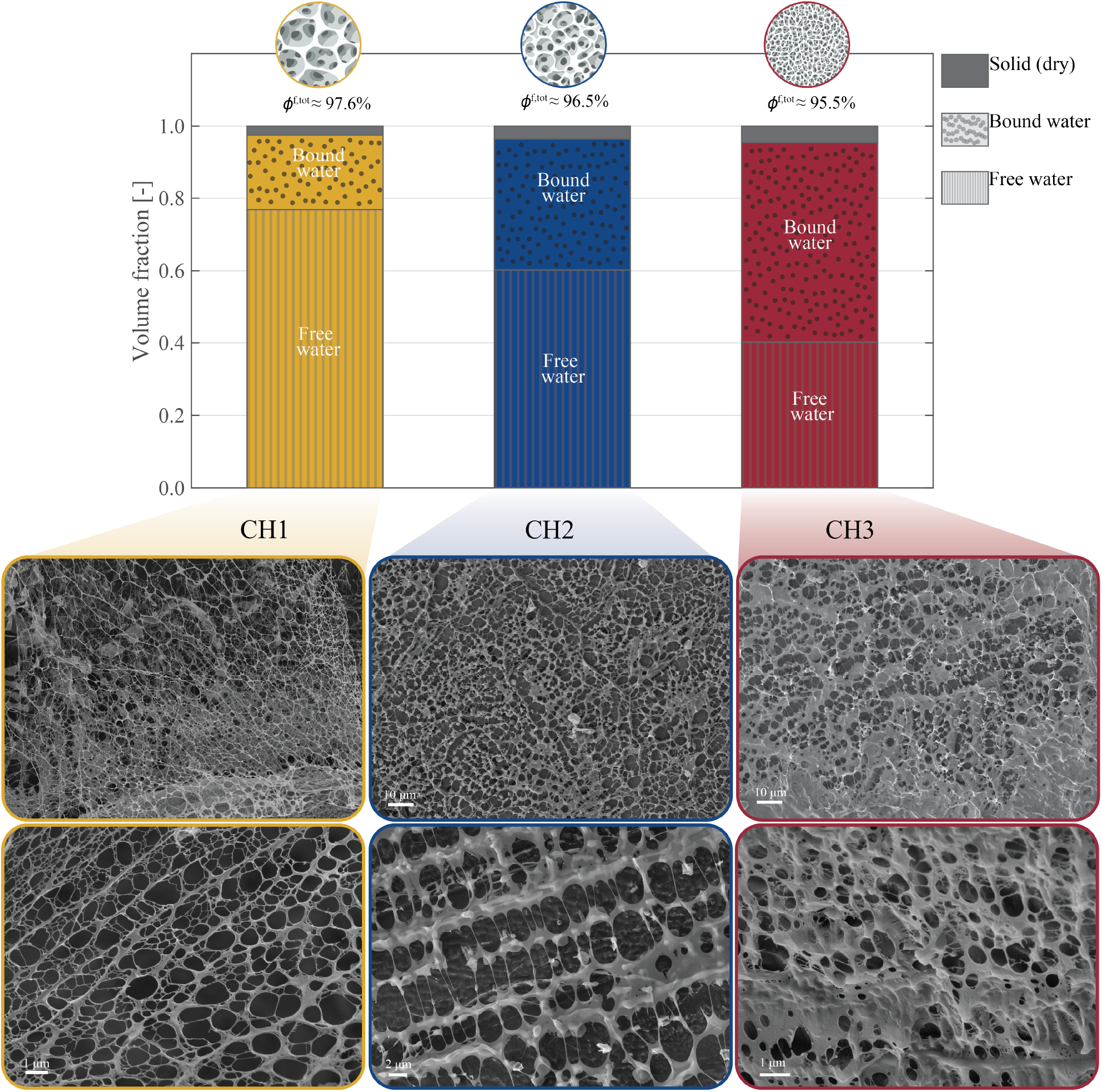
Summary of the identified fluid volume fractions in the three hydrogel compositions (median values). The parameter ‘Free water’, refers to the amount of free-flowing fluid that was mechanically displaced using the incremental consolidation protocol, identified with the effective porosity 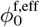. The parameter ‘Bound water’ is a derived quantity calculated from the difference between the total fluid volume fraction *ϕ*^f,tot^, obtained from dehydration and gravimetric analysis, and the mechanically displaced water volume. It represents the portion of the total fluid volume that remains bound within the sample even after compaction. The fraction of the polymer remaining after full dehydration is referred to as ‘Solid (dry)’. At the bottom, representative scanning electron microscopy images of each hydrogel composition in the reference swollen state are shown.

Representative SEM micrographs for each composition are shown in Fig. 8. The median values of the effective porosity, along with their IQR, are shown in Fig. 9(a). Pairwise comparisons were conducted using the two-sided Mann– Whitney *U* test (unadjusted *p*-values). All pairwise differences were highly significant: CH1 vs CH2 (*p* ≈8.23 ×10^−5^), CH1 vs CH3 (*p* ≈4.57 ×10^−5^), and CH2 vs CH3 (*p* ≈2.80 ×10^−4^). In all cases, Cliff’s *δ* equals 1.0, indicating complete separation between groups. The polymeric (solid) phases in the three compositions accounted for 2.3%, 3.4%, and 4.6% of the total mass, respectively. These results confirm that higher polymer concentrations correspond to lower free-water fractions. Consequently, the amount of free water is directly tunable through the polymer concentration, which is crucial for the design of mechano-responsive soft devices and related applications. A detailed graphical representation, together with the relative roles of PVA and PHY, is provided in the Supplementary Material. The extreme compaction is achieved through a drastic reduction in porosity, as shown by the comparison of representative images of the swollen state with those of the compacted state reported in Fig. 10 below. However, potential damage to the interconnected porous network cannot be completely ruled out, which may affect the computation of the effective porosity in the reference state 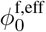.

**Figure 9:**
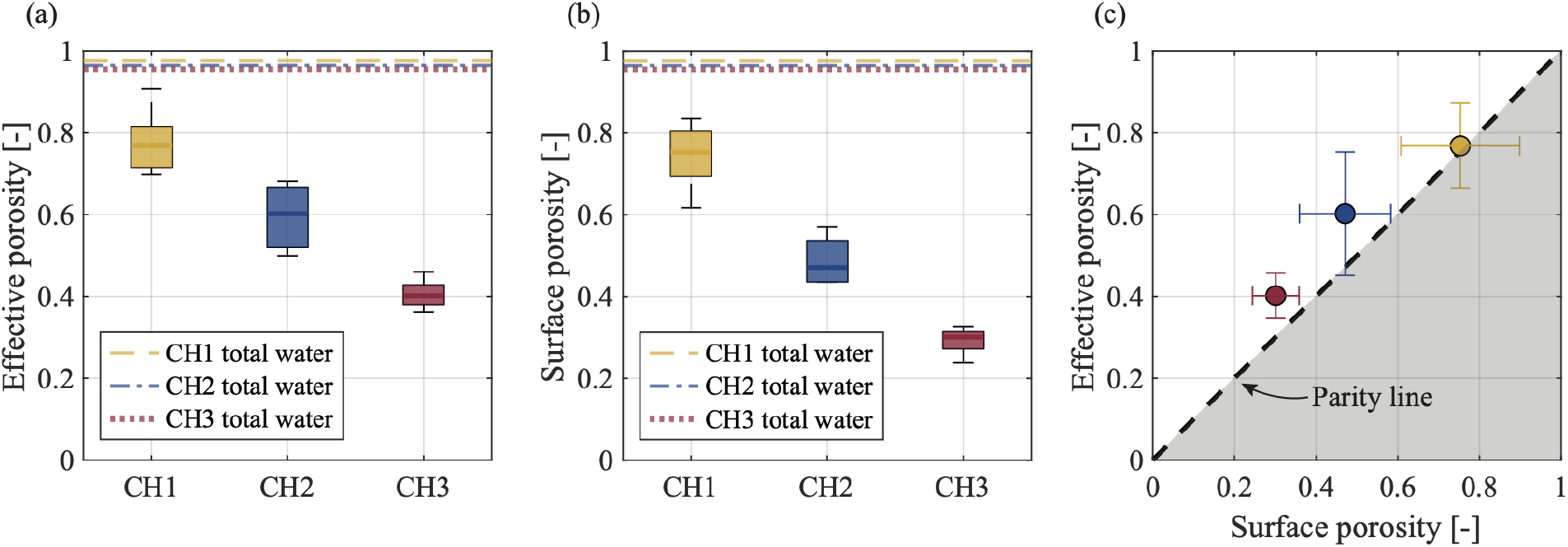
Effective and surface porosity: (a) effective porosity as amount of free water quantified using the incremental consolidation method (median and IQR); (b) surface porosity derived from the cryo-SEM analysis (median and IQR); (c) comparison between effective and surface porosity. Data are shown as media and IQR.

**Figure 10:**
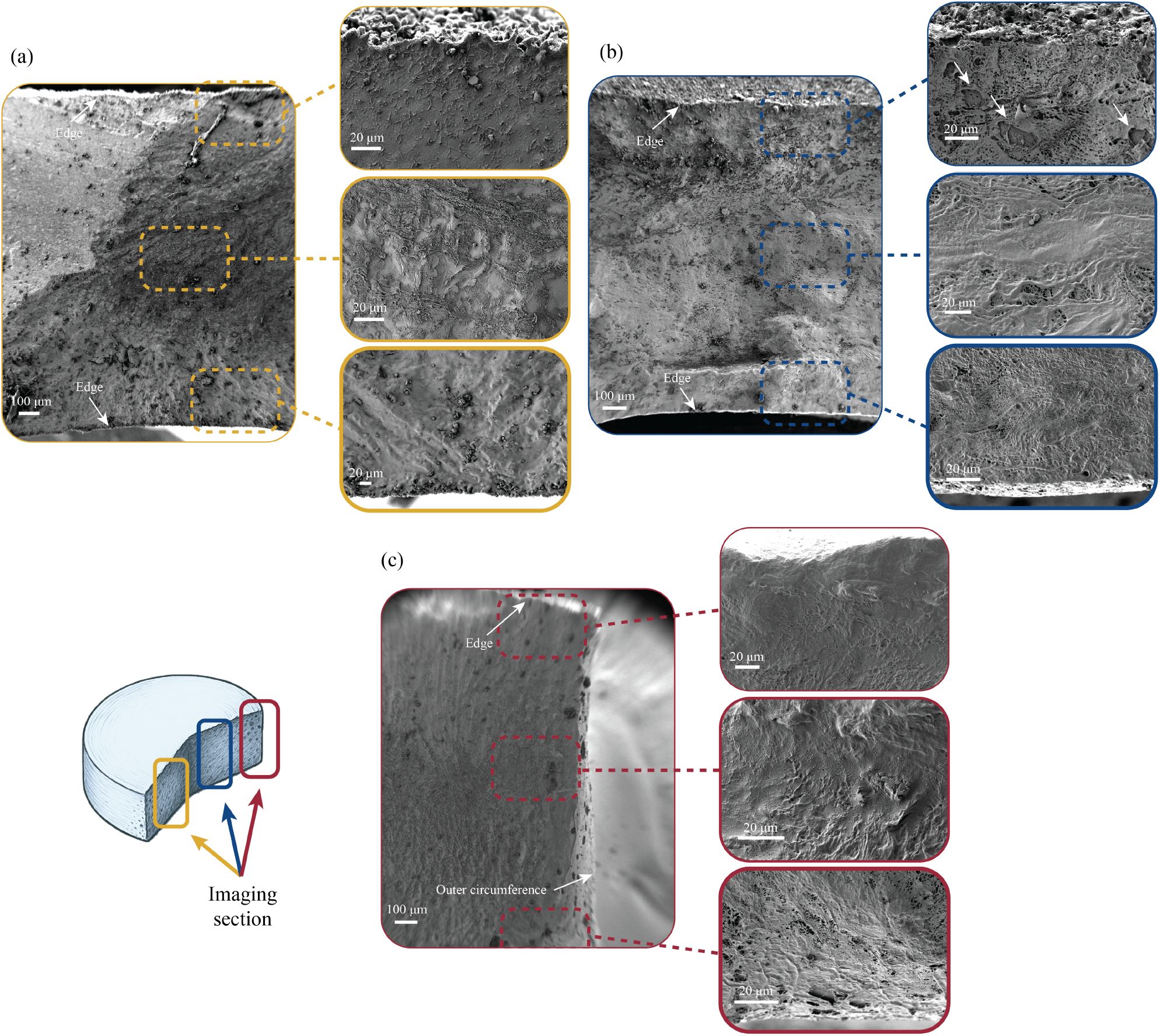
Scanning electron microscopy images of compacted hydrogels: (a) CH1, (b) CH2 and (c) CH3. Images show a front view from top to bottom edge and magnifications of the areas around the top edge, the center area, and the area around the bottom edge.

### 4.4 Cryo-SEM analyses: surface porosity and its relation to effective porosity

Using identically prepared cryo-SEM sections, we derived a quantitative metric of surface porosity for the composite hydrogels, defined as the fraction of pore space within a reference area. Further details on the methods are provided in Sec. 3.4.1, while median values and IQR are shown in Fig. 9(b). All pairwise comparisons between hydrogel compositions were found statistically significant (two-sided Mann–Whitney *U* test; unadjusted *p*-values): CH1 vs CH2 (*p* ≈1.19 ×10^−2^), CH1 vs CH3 (*p* ≈7.94 ×10^−3^), and CH2 vs CH3 (*p* ≈1.19 ×10^−2^). Cliff’s *δ* for each SEM pair is 1.0, indicating complete separation between the groups of the image-based porosity metric. Furthermore, we compared the surface porosity derived from the cryo-SEM method with the effective porosity calculated from the incremental consolidation experiments. The mechanical measurement represents a macroscopic, coarse-grained scale that is representative of the effective pore connectivity, as it captures the gross fluid transport through the polymer matrix that static, localized images cannot resolve. For CH1, the two measures closely agree: the difference is negligible (about 2%), not statistically significant (*p* ≈ 0.83) and the effect size is small (Cliff’s *δ* ≈ 0.10), indicating no practical discrepancy between surface and mechanical estimates for the lowest-polymer composition. For CH2, the mechanically-derived effective porosity exceeds the surface estimate by approximately 22%. Although this difference did not reach the conventional significance threshold (*p* ≈0.08), the effect size is moderate (Cliff’s *δ*≈ 0.60), suggesting a consistent directional effect that may be sufficiently pronounced in the present sample. Finally, CH3 shows a pronounced discrepancy: the effective porosity is substantially higher (25%) than the surface-derived value, the difference is statistically significant (*p*≪ 0.01), and the effect size is very large (Cliff’s *δ* ≈1.0). Taken together, these results suggest that surface imaging underestimates the fraction of free water as polymer content rises. A direct visual comparison is provided in Fig. 9(c). The parity line marks perfect agreement between methods; points above this line indicate higher surface-porosity estimates, while points below (gray-shaded region) indicate higher mechanically-determined values.

We hypothesize two effects contributing to the increasing mismatch between image-derived surface porosity and mechanically measured effective porosity with rising polymer content. First, in hydrogels with higher polymer concentration, the cryo-SEM preparation and freeze-drying can cause stronger mechanical distortions, local collapse or rearrangement of the delicate structure, altering surface metrics relative to the hydrated state [37, 103, 104]. The potential introduction of structural artifacts following cryogenic treatment – particularly in soft and highly hydrated materials – remains a subject of significant debate and could be evaluated through comparison with additional microscopy methods [45]. Second, since surface porosity is quantified in the undeformed state, it captures only the area fraction of open voids after sublimation and cannot account for water retained within the swollen polymer mesh. In contrast, the incremental consolidation experiment expels water under load and, over the relaxation period, allows water to migrate out of the polymer mesh (intra-mesh water), which is thus mechanically released by deformation. The growing mismatch with polymer content is therefore consistent with (i) stronger polymer–water interactions and potential distortions, and (ii) an increasing reservoir of intra–mesh water that is invisible to surface analysis but is mobilized during consolidation. Taken together, these observations support our claim that the consolidation-based approach allows for a practical estimation of the effective porosity, especially with regard to its mechanical importance for the modeling of biphasic materials.

### 4.5 Cryo-SEM analyses: morphology of compacted hydrogels and limitations of confined compression

Figure 10 shows cryo-SEM representative images of the hydrogels following completion of the incremental consolidation protocol. Hydrogel composites CH1 and CH3 (Fig. 10(a),(c)) appear fully and uniformly compacted, whereas CH2 (Fig. 10(b)) shows some open pores near the top surface, indicated by arrows. This qualitative observation can have relevant implications for the analysis of the incremental consolidation experiment.

During confined compression-consolidation, pore fluid is driven toward the rigid, porous bottom, leading to a pressure gradient and inhomogeneous fluid flow in the sample. Regions near to the drained boundary lose fluid first and the local solid matrix compacts, while regions further away remain more fluid-filled and compliant. The resulting state is therefore characterized by a gradient in porosity, from the drained side to the opposite boundary, rather than a spatially uniform distribution. The practical consequence is that boundary-driven, asymmetric compaction could influence the estimates of the free-water fraction and thus of the effective porosity. To mitigate these effects, the following strategies could be, e.g., adopting longer equilibration times, and/or finer resolved compression steps.

### 4.6 Cryo-SEM analyses: scale-dependence of surface porosity

In the cryo-SEM analysis, hydrogel samples were imaged at multiple magnifications and cryo-fractionated under controlled atmosphere to expose the bulk microstructure and minimize preparation artifacts. Although all samples appeared macroscopically homogeneous across the different tests and experiments, PVA- and polysaccharide-based hydrogels produced by freeze–thawing can display microstructural inhomogeneity at scales larger than a few micrometers. Associated data and images have been reported in studies using only PVA-based hydrogels [46, 105, 106] as well as with various PVA-composites [34, 35, 107]. To mitigate the influence of microscale inhomogeneous pore space, we (i) analyzed multiple images from multiple samples and (ii) investigated the scale dependence of the derived metric. The dependence of surface porosity on structure size and area fraction is shown in Fig. 11 for three representative samples. Figures 11(a)-(c) show the derived surface porosity as a function of structure size, expressed as the side length *L* of the moving window used in the analysis (Sec. 3.4.1). Regardless of the composition of the hydrogel, all curves converge to a constant value. For CH1, the curve begins at 100% and then flattens, whereas for CH2 and CH3, the curves start at 0% surface porosity. This behavior arises because, at structure sizes close to zero, the window evaluates individual pixels: after binarization, each pixel is either pore space or solid, corresponding to a 100% or 0% porosity, respectively. At small window sizes, the data spread represented as IQR is comparatively large, reflecting microscopic inhomogeneities that become visible at small scales. As the structure size increases, the spread decreases and the IQRs narrow, indicating that at these scales microscopic inhomogeneities have a negligible influence and the derived metric can be considered representative. A similar trend appears when porosity is plotted as a function of normalized window area (area fraction). For the analyzed image sizes, the median values therefore capture the central tendency of the underlying local-porosity distribution. The dashed vertical lines mark the area defined as representative area element (RAE). The RAE is determined based on a user-set tolerance (±5% of the converged median value). Therefore, a RAE with the corresponding physical structure size can be considered as a threshold for deriving the surface porosity. Although cryogenic methods remain state-of-the-art method for pore morphology and pore size analysis in soft, highly hydrated hydrogels [103, 108, 109], we have to mention some limitations [40]. Despite using rapid freezing methods to minimize artifacts, the formation of ice crystals can distort the microstructure [109], and electrical charging during the scan can alter the acquired images [110]. It should thus be emphasized that the resulting structures must be interpreted accordingly.

**Figure 11:**
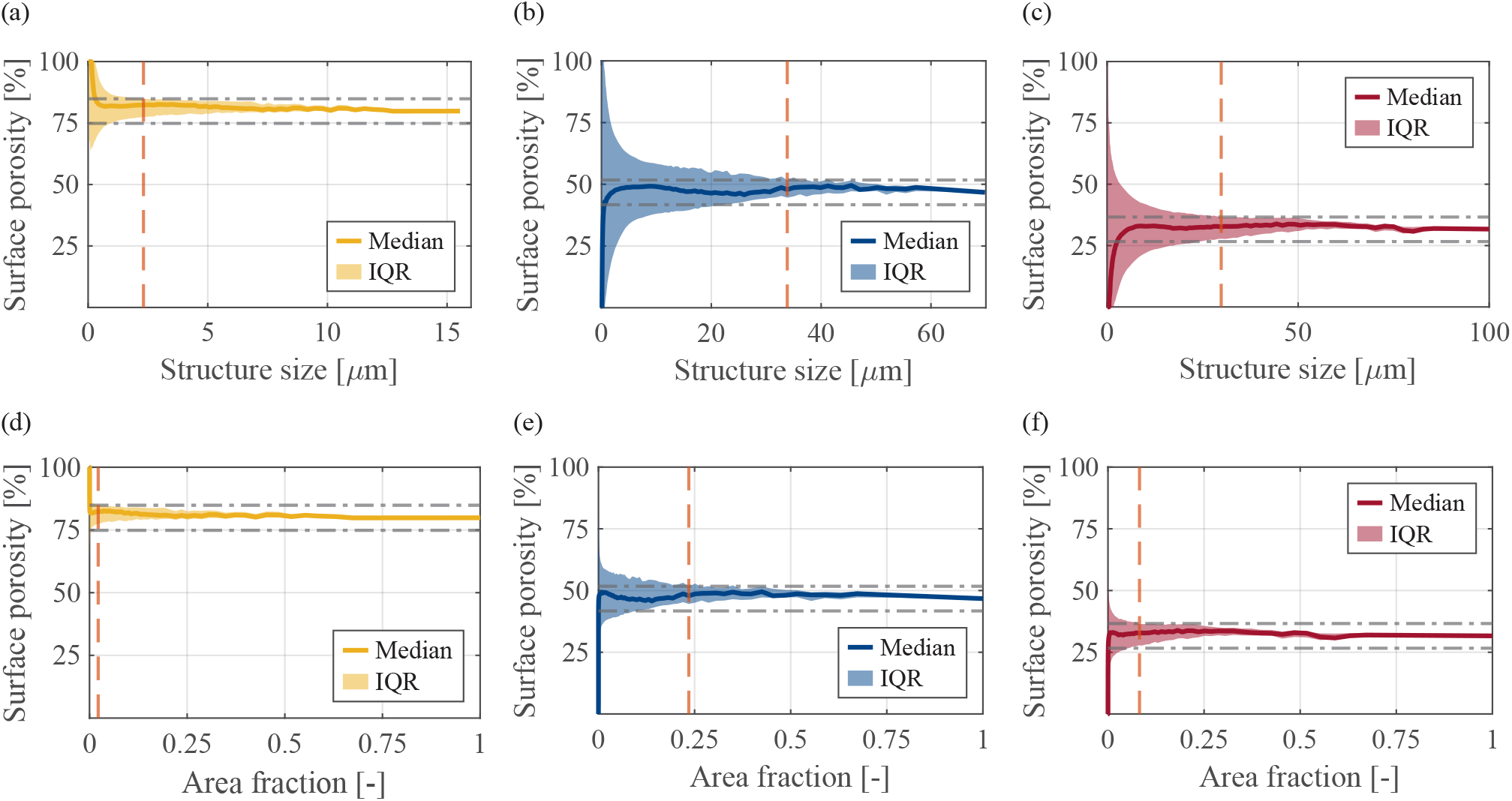
Scale- and size dependence of surface porosity, for representative compositions of CH1, CH2 and CH3 hydrogels: (a)-(c) porosity over the structure size, i.e. the side length of an analyzed scanning electron microscopy image; (d)-(f) same data referenced to the image area fraction. Dashed, horizontal lines visualize a region of ±5% of the converged value. Dashed, vertical lines indicate where the IQR range exceeds the user-defined 5%-range and can be used to identify a representative area element.

## 5 Conclusion

The amount and state of water in hydrogels decisively determine their mechanical and transport properties, and thus their suitability for different applications. Using three different ultra-soft, freeze-thawed composite hydrogels, we were able to demonstrate that incremental consolidation is a reproducible method to compact the hydrogel matrix and directly derive the mechanically releasable free-water content. By quantifying the displaced fluid volume during controlled consolidation steps, an effective porosity for moderate loading scenarios was derived that can be directly used in the modeling of biphasic materials. Importantly, it was found that the effective porosity scales linearly with the polymer concentration. This suggests that the free-water content can be precisely controlled by the formulation while keeping other parameters constant. The surface porosity determined by cryo-SEM analysis consistently yielded lower values than the mechanically determined free-water fraction, showing increasing deviation with increasing polymer concentrations. By contrast, the total water fraction obtained from the gravimetric analysis includes a large fraction of fluid which cannot be mechanically mobilized.

Our finding that a purely mechanical protocol can provide an estimate of the releasable free-water fraction has two immediate, complementary implications:

i. For macroporous soft hydrogels based on PVA and produced via phase-templating, the proposed protocol enables the direct, experimental determination of the effective porosity. In addition, the steady-state points at prescribed stretches and associated equilibrium stresses can be used directly to calibrate the material behavior of the solid phase.
ii. The three different hydrogel formulations allow for controlled tuning of the free-water fraction that can be released under moderate loadings. We report the relevant loading parameters (up to 2 N, area-normalized to 25 kPa, macroscopic stretches 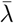 up to ≈0.4–0.8, depending on the composition). For applications based on mechano-triggered fluid release, e.g., controlled fluid delivery, responsive scaffolds, or surface-conforming interfaces where fluid transport is mechanically modulated, the free-water fraction together with the reported load/stretch thresholds are crucial design parameters. Knowledge of both the amount of releasable fluid and the mechanical stimulus required for mobilization enables the engineering of hydrogel systems with predictable release kinetics and trigger thresholds.

In practice, this means that the protocol can be used both to parameterize mechanical simulations for predictive studies and to support material selection for applications where knowledge and control of mechanically driven fluid transport are required. Potential next steps in this context would be to quantitatively compare the fractions using spectroscopic methods and thermal analysis (e.g., DSC) that can specifically classify binding states of water, as well as an investigation of temperature-dependent behavior. Furthermore, the temperature and energy input associated with very high mechanical loads might facilitate the detachment of bound water through mechanisms such as internal friction or localized heat generation. Although these contributions are considered minor compared to the large displaced volumes of free water characterized here, they could be of specific interest for the release kinetics of encapsulated species under mechanical stimulus, as the mobilization of the bound layer could play a crucial role.

## 6 Acknowledgments

This research was funded in whole or in part by the Austrian Science Fund (FWF) 10.55776/I4828. For the purpose of open access, the author has applied a CC BY public copyright license to any author accepted manuscript version arising from this submission. We are grateful to Kerstin Hingerl and Philip Schulner for helping with the imaging and the experiments, and to Maximilian Wollner for the support and inspiring discussions. Figures, images, and artwork, including the graphical abstract, were created in whole or in part using BioRender.com.

## A Fundamentals of Mixture Theories

### A.1 Kinematics

The theory of mixtures is a continuum approach to describing the mechanical behavior of materials with multiple phases and constituents. A mixture can be viewed as a sequence of bodies that can simultaneously occupy the same region of space. We begin by defining the motion of phase *α*, i.e.

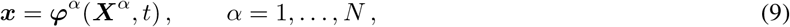

which relates each particle located at ***X***^*α*^ in the reference configuration ℬ^*α*^ of the *α*-th phase to the spatial position ***x*** at time *t* in the current configuration of the mixture ℬ_*t*_.

The deformation gradient associated to this motion is defined by

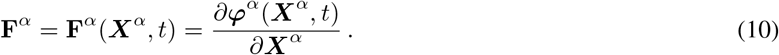

The velocity of phase *α* is defined as

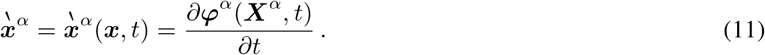

### A.2 Balance laws

The fundamental axioms of mass and momentum balance are formulated in a spatial setting for an infinitesimal volume d*v* attached to ***x*** ∈ ℬ_*t*_, which is called the volume element of the mixture. In the absence of mass exchange between the phases, the mass balances are written as

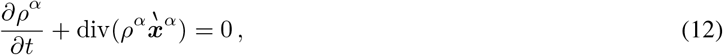

where *ρ*^*α*^ is the apparent density of phase *α*. Physically, it corresponds to the mass of phase *α* per unit volume of the mixture. For immiscible mixtures, in which each phase occupies a volume separate from all others, the true density 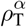 can also be defined as the mass of phase *α* per unit volume that this phase occupies in the mixture. The apparent and true densities are related through the volume fraction *ϕ*^*α*^, which is defined as follows:

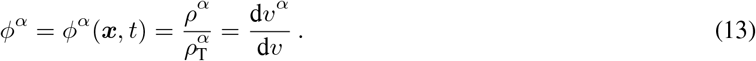

Neglecting dynamic effects and gravity, the linear momentum balance for phase *α* is given by

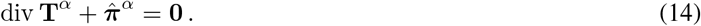

In Eq. (14), **T**^*α*^ is the partial or apparent Cauchy stress tensor of the phase *α*, defined such that the force vector acting on the *α*-th phase per unit area of the mixture satisfies ***t***^*α*^ = ***n***· **T**^*α*^, with ***n*** the unit normal vector. The vector 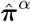 accounts for the exchange of linear momentum between phase *α* and the rest of the mixture, satisfying 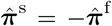 in a biphasic mixture.

An interesting special case concerns saturated mixtures of incompressible phases. In this case, the volume fractions are subject to the constraint ∑_*α*_ *ϕ*^*α*^ = 1, and the true density 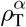 is constant. Therefore, the following special cases of the mass balance, i.e. (12), for the single phase and the mixture arise. Thus,

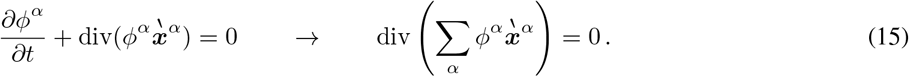

The constraint on the mixture expressed by (15) generates a work-less reaction force which physically corresponds to an interface pressure required to maintain contact between the phases.

### A.3 Uniaxial boundary-value problem

The preceding equations can be used to formulate a BVP for a biphasic porous medium under isothermal conditions. The unknown fields include the motion functions ***φ***^*α*^, the volume fractions *ϕ*^*α*^, the stress tensors **T**^*α*^, and the exchange terms 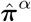. The solution of the complete BVP is not considered in this work, as the focus is exclusively on the steady state.

From Eq. (15), with the assumption of mixture saturation, we derive the following constraint, i.e.

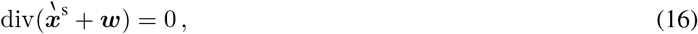

where 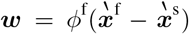 is known as discharge velocity. Constraint (16) is classically enforced through a Lagrange multiplier *p*, which introduces an additional unknown identifiable as the interstitial fluid pressure [76].

In the context of the uniaxial BVP presented in Sec. 2.1 and illustrated in Fig. 2(b), for an isotropic solid and an inviscid fluid, Eq. (14) and Eq. (16) reduce to

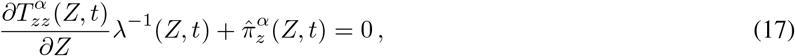

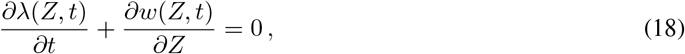

where we made use of the identity ∂(•)*/*∂*Z* = *λ*∂(•)*/*∂*z*. Here, *λ* defines the generally inhomogeneous stretch ratio in the longitudinal direction, i.e., *λ* = *λ*(*Z, t*) = ∂*φ*^s^ (*Z, t*)*/*∂*Z*. With the boundary conditions

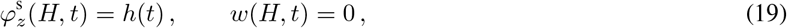

we can integrate Eq. (18) to obtain

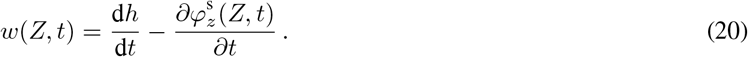

We define the steady state as the condition in which both the solid and fluid are stationary. Consequently, with *w*(*Z, t*^eq^) = 0 the continuity equation (18) shows that the stretch is constant in time. Furthermore, the momentum exchange term in Eq. (17) vanishes, leaving the two balances of linear momentum to be trivially satisfied by a spatially homogeneous stretch. Therefore, the Cauchy stress tensor of the solid at equilibrium reads

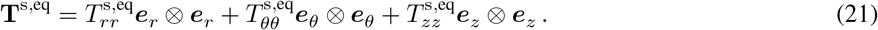

## Notes

### Competing Interest Statement

The authors have declared no competing interest.

## References

[1] Antonio Elia Forte et al. “A composite hydrogel for brain tissue phantoms”. In: Materials & Design 112 (2016), pp. 227–238. DOI: 10.1016/j.matdes.2016.09.063.

[2] Peilin Lu et al. “Harnessing the Potential of Hydrogels for Advanced Therapeutic Applications: current Achievements and Future Directions”. In: Signal Transduction and Targeted Therapy 9.1 (2024), p. 166. ISSN: 2059-3635. DOI: 10.1038/s41392-024-01852-x.

[3] Roberto Brighenti, Mattia P. Cosma, and Noy Cohen. “Mechanics and Physics of the Light-Driven Response of Hydrogels”. In: Mechanics Research Communications 129 (2023), p. 104077. ISSN: 00936413. DOI: 10.1016/j.mechrescom.2023.104077.

[4] Hussein M. El-Husseiny et al. “Smart/stimuli-responsive hydrogels: Cutting-edge platforms for tissue engineering and other biomedical applications”. In: Materials Today Bio 13 (2022), p. 100186. ISSN: 2590-0064. DOI: 10.1016/j.mtbio.2021.100186.

[5] Simin Cheng, Ruiqi Zhu, and Xiaomin Xu. “Hydrogels for next Generation Neural Interfaces”. In: Communications Materials 5.1 (2024), p. 99. ISSN: 2662-4443. DOI: 10.1038/s43246-024-00541-0.

[6] Ling Wang et al. “Biomimetic Design and Integrated Biofabrication of an In-Vitro Three-Dimensional Multi-Scale Multilayer Cortical Model”. In: Materials Today Bio 28 (2024), p. 101176. ISSN: 25900064. DOI: 10.1016/j.mtbio.2024.101176.

[7] Cunjiang Yu et al. “Electronically Programmable, Reversible Shape Change in Two- and Three-Dimensional Hydrogel Structures”. In: Advanced Materials 25.11 (2013), pp. 1541–1546. ISSN: 0935-9648, 1521-4095. DOI: 10.1002/adma.201204180.

[8] Ottavia Bettucci, Giovanni Maria Matrone, and Francesca Santoro. “Conductive Polymer-Based Bioelectronic Platforms toward Sustainable and Biointegrated Devices: a Journey from Skin to Brain across Human Body Interfaces”. In: Advanced Materials Technologies 7.2 (2022), p. 2100293. ISSN: 2365-709X, 2365-709X. DOI: 10.1002/admt.202100293.

[9] Xiao Wang et al. “Bioadhesive and Conductive Hydrogel-Integrated Brain-Machine Interfaces for Conformal and Immune-Evasive Contact with Brain Tissue”. In: Matter 5.4 (2022), pp. 1204–1223. ISSN: 25902385. DOI: 10.1016/j.matt.2022.01.012.

[10] N. A. Peppas et al. “Physicochemical Foundations and Structural Design of Hydrogels in Medicine and Biology”. In: Annual Review of Biomedical Engineering 2.1 (2000), pp. 9–29. ISSN: 1523-9829, 1545-4274. DOI: 10.1146/annurev.bioeng.2.1.9.

[11] Masao Doi. “Gel Dynamics”. In: Journal of the Physical Society of Japan 78.5 (2009), p. 052001. ISSN: 0031-9015. DOI: 10.1143/JPSJ.78.052001.

[12] Vladimir M. Gun’ko, Irina N. Savina, and Sergey V. Mikhalovsky. “Properties of Water Bound in Hydrogels”. In: Gels 3.4 (2017). ISSN: 2310-2861. DOI: 10.3390/gels3040037.

[13] Yue Yuan et al. “Water: the Soul of Hydrogels”. In: Progress in Materials Science 148 (2025), p. 101378. DOI: 10.1016/j.pmatsci.2024.101378.

[14] Lifeng Li et al. “Effect of Water State and Polymer Chain Motion on the Mechanical Properties of a Bacterial Cellulose and Polyvinyl Alcohol (BC/PVA) Hydrogel”. In: RSC Advances 5.32 (2015), pp. 25525–25531. ISSN: 2046-2069. DOI: 10.1039/C4RA11594E.

[15] Allan S Hoffman. “Hydrogels for Biomedical Applications”. In: Advanced Drug Delivery Reviews 64 (2012), pp. 18–23. ISSN: 0169409X. DOI: 10.1016/j.addr.2012.09.010.

[16] Jamie L. Hernandez and Kim A. Woodrow. “Medical Applications of Porous Biomaterials: features of Porosity and Tissue-Specific Implications for Biocompatibility”. In: Advanced Healthcare Materials 11.9 (2022), p. 2102087. ISSN: 2192-2640. DOI: 10.1002/adhm.202102087.

[17] M. A. Kristine Tolentino et al. “Decoding Hydrogel Porosity: advancing the Structural Analysis of Hydrogels for Biomedical Applications”. In: Advanced Healthcare Materials 14.22 (2025), p. 2500658. ISSN: 2192-2640, 2192-2659. DOI: 10.1002/adhm.202500658.

[18] Chuan Li et al. “Fine Tuning Water States in Hydrogels for High Voltage Aqueous Batteries”. In: ACS Nano 18.4 (2024), pp. 3101–3114. ISSN: 1936-0851. DOI: 10.1021/acsnano.3c08398.

[19] Francis X. Quinn et al. “Water in Hydrogels. 1. A Study of Water in Poly(N-vinyl-2-pyrrolidone/Methyl Methacrylate) Copolymer”. In: Macromolecules 21.11 (1988), pp. 3191–3198. ISSN: 0024-9297, 1520-5835. DOI: 10.1021/ma00189a012.

[20] Reza Foudazi et al. “Porous Hydrogels: present Challenges and Future Opportunities”. In: Langmuir 39.6 (2023), pp. 2092–2111. ISSN: 0743-7463. DOI: 10.1021/acs.langmuir.2c02253.

[21] Gustav Graeber et al. “Intrinsic Water Transport in Moisture-Capturing Hydrogels”. In: Nano Letters 24.13 (2024), pp. 3858–3865. ISSN: 1530-6984, 1530-6992. DOI: 10.1021/acs.nanolett.3c04191.

[22] Clifford Truesdell. Rational Thermodynamics. New York: McGraw Hill, 1969.

[23] Jacob Bear. Dynamics of Fluids in Porous Media. New York: Elsevier, 1972.

[24] Ray M. Bowen. “Theory of Mixtures”. In: Continuum Physics. Elsevier, 1976, pp. 1–127. ISBN: 978-0-12-240803-8. DOI: 10.1016/B978-0-12-240803-8.50017-7.

[25] Reint De Boer. Theory of Porous Media. Berlin, Heidelberg: Springer Berlin Heidelberg, 2000.

[26] Olivier Coussy. Poromechanics. Chichester: Wiley, 2003. DOI: 10.1002/0470092718.

[27] Mark H. Holmes and V.C. Mow. “The Nonlinear Characteristics of Soft Gels and Hydrated Connective Tissues in Ultrafiltration”. In: Journal of Biomechanics 23.11 (1990), pp. 1145–1156. ISSN: 00219290. DOI: 10.1016/0021-9290(90)90007-P.

[28] Michael A. Soltz and Gerard A. Ateshian. “Experimental Verification and Theoretical Prediction of Cartilage Interstitial Fluid Pressurization at an Impermeable Contact Interface in Confined Compression”. In: Journal of Biomechanics 31.10 (1998), pp. 927–934. ISSN: 00219290. DOI: 10.1016/S0021-9290(98)00105-5.

[29] LRG Treloar. The Physics of Rubber Elasticity. New York: Oxford University Press, 1975. ISBN: 978-0-19-152330-4.

[30] Alexander E. Ehret et al. “Inverse Poroelasticity as a Fundamental Mechanism in Biomechanics and Mechanobiology”. In: Nature Communications 8.1 (2017), p. 1002. ISSN: 2041-1723. DOI: 10.1038/s41467-017-00801-3.

[31] Krzysztof M. Graczyk and Maciej Matyka. “Predicting Porosity, Permeability, and Tortuosity of Porous Media from Images by Deep Learning”. In: Scientific Reports 10.1 (2020), p. 21488. ISSN: 2045-2322. DOI: 10.1038/s41598-020-78415-x.

[32] Malak Alaa Eddine et al. “Large and Nonlinear Permeability Amplification with Polymeric Additives in Hydrogel Membranes”. In: Macromolecules 55.21 (2022), pp. 9841–9850. ISSN: 0024-9297, 1520-5835. DOI: 10.1021/acs.macromol.2c01462.

[33] Anuj Kumar and Sung Soo Han. “PVA-based Hydrogels for Tissue Engineering: a Review”. In: International Journal of Polymeric Materials and Polymeric Biomaterials 66.4 (2017), pp. 159–182. ISSN: 0091-4037. DOI: 10.1080/00914037.2016.1190930.

[34] Weichang Li et al. “Preparation and Characterization of PVA-PEEK/PVA-β-TCP Bilayered Hydrogels for Articular Cartilage Tissue Repair”. In: Composites Science and Technology 128 (2016), pp. 58–64. ISSN: 02663538. DOI: 10.1016/j.compscitech.2016.03.013.

[35] Liang Ma et al. “A High-Water Retention, Self-Healing Hydrogel Thyroid Model for Surgical Training”. In: Materials Today Bio 29 (2024), p. 101334. ISSN: 25900064. DOI: 10.1016/j.mtbio.2024.101334.

[36] Christie M. Hassan and Nikolaos A. Peppas. “Structure and Morphology of Freeze/Thawed PVA Hydrogels”. In: Macromolecules 33.7 (2000), pp. 2472–2479. ISSN: 0024-9297, 1520-5835. DOI: 10.1021/ma9907587.

[37] Jérôme Grenier et al. “Mechanisms of Pore Formation in Hydrogel Scaffolds Textured by Freeze-Drying”. In: Acta Biomaterialia 94 (2019), pp. 195–203. ISSN: 17427061. DOI: 10.1016/j.actbio.2019.05.070.

[38] Sitao Wang et al. “Porosity Engineering of Dried Smart Poly(N -Isopropylacrylamide) Hydrogels for Gas Sensing”. In: Biomacromolecules 25.5 (2024), pp. 2715–2727. ISSN: 1525-7797, 1526-4602. DOI: 10.1021/acs.biomac.3c00738.

[39] Todd M. Alam et al. “Characterization of Free, Restricted, and Entrapped Water Environments in Poly(N-isopropyl Acrylamide) Hydrogels via 1H HRMAS PFG NMR Spectroscopy”. In: Journal of Polymer Science Part B: Polymer Physics 52.23 (2014), pp. 1521–1527. ISSN: 0887-6266, 1099-0488. DOI: 10.1002/polb.23591.

[40] Martin Bartoš, Tomáš Suchý, and René Foltán. “Note on the Use of Different Approaches to Determine the Pore Sizes of Tissue Engineering Scaffolds: what Do We Measure?” In: BioMedical Engineering OnLine 17.1 (2018), p. 110. ISSN: 1475-925X. DOI: 10.1186/s12938-018-0543-z.

[41] Miroslava Dušková-Smrčková et al. “Communicating Macropores in PHEMA-based Hydrogels for Cell Seeding: Probabilistic Open Pore Simulation and Direct Micro-CT Proof”. In: Materials & Design 198 (2021), p. 109312. ISSN: 02641275. DOI: 10.1016/j.matdes.2020.109312.

[42] Nilly Hojat et al. “Automatic Pore Size Measurements from Scanning Electron Microscopy Images of Porous Scaffolds”. In: Journal of Porous Materials 30.1 (2023), pp. 93–101. ISSN: 1380-2224, 1573-4854. DOI: 10.1007/s10934-022-01309-y.

[43] Rasoul Mirghafari, Daniel Bell, and Olga Barrera. “An image-based approach for advanced statistical quantification of architectural parameters and permeability in porous biological media”. In: Acta Biomaterialia 203 (2025), pp. 509–522. DOI: 10.1016/j.actbio.2025.06.022.

[44] Isadora C. Carvalho and Herman S. Mansur. “Engineered 3D-scaffolds of Photocrosslinked Chitosan-Gelatin Hydrogel Hybrids for Chronic Wound Dressings and Regeneration”. In: Materials Science and Engineering: C 78 (2017), pp. 690–705. ISSN: 09284931. DOI: 10.1016/j.msec.2017.04.126.

[45] Zhansaya Kaberova et al. “Microscopic Structure of Swollen Hydrogels by Scanning Electron and Light Microscopies: Artifacts and Reality”. In: Polymers 12.3 (2020), p. 578. ISSN: 2073-4360. DOI: 10.3390/polym12030578.

[46] Rosangela Mastrangelo et al. “‘Twin-Chain’ Hydrogels with Tailored Porosity, Surface Roughness, and Cleaning Capabilities”. In: Langmuir 41.16 (2025), pp. 10238–10249. ISSN: 0743-7463, 1520-5827. DOI: 10.1021/acs.langmuir.4c05381.

[47] Masanobu Nagura et al. “States of Water in Poly(Vinyl Alcohol) Hydrogels”. In: Polymer Gels and Networks 5.5 (1997), pp. 455–468. ISSN: 09667822. DOI: 10.1016/S0966-7822(97)00016-6.

[48] Chun-Wei Chang et al. “An Investigation of Water Status in Gelatin Methacrylate Hydrogels by Means of Water Relaxometry and Differential Scanning Calorimetry”. In: Journal of Materials Chemistry B 12.26 (2024), pp. 6328–6341. ISSN: 2050-750X, 2050-7518. DOI: 10.1039/D4TB00053F.

[49] Jirong Yang et al. “Influence of Hydrogel Network Microstructures on Mesenchymal Stem Cell Chondrogenesis in Vitro and in Vivo”. In: Acta Biomaterialia 91 (2019), pp. 159–172. ISSN: 17427061. DOI: 10.1016/j.actbio.2019.04.054.

[50] Louise Griveau et al. “Design and characterization of an in vivo injectable hydrogel with effervescently generated porosity for regenerative medicine applications”. In: Acta Biomaterialia 140 (2022), pp. 324–337. ISSN: 1742-7061. DOI: 10.1016/j.actbio.2021.11.036.

[51] Monica Jane Emerson et al. “Statistical Validation of Individual Fibre Segmentation from Tomograms and Microscopy”. In: Composites Science and Technology 160 (2018), pp. 208–215. ISSN: 02663538. DOI: 10.1016/j.compscitech.2018.03.027.

[52] W.Y. Gu et al. “New Insight into Deformation-Dependent Hydraulic Permeability of Gels and Cartilage, and Dynamic Behavior of Agarose Gels in Confined Compression”. In: Journal of Biomechanics 36.4 (2003), pp. 593– 598. ISSN: 00219290. DOI: 10.1016/S0021-9290(02)00437-2.

[53] Jonathan A. Kluge et al. “The Consolidation Behavior of Silk Hydrogels”. In: Journal of the Mechanical Behavior of Biomedical Materials 3.3 (2010), pp. 278–289. ISSN: 17516161. DOI: 10.1016/j.jmbbm.2009.12.001.

[54] Qunli Liu, Ghatu Subhash, and David F. Moore. “Loading Velocity Dependent Permeability in Agarose Gel under Compression”. In: Journal of the Mechanical Behavior of Biomedical Materials 4.7 (2011), pp. 974–982. ISSN: 17516161. DOI: 10.1016/j.jmbbm.2011.02.009.

[55] Reinder W. Roos, Rob Petterson, and Jacques M. Huyghe. “Confined Compression and Torsion Experiments on a pHEMA Gel in Various Bath Concentrations”. In: Biomechanics and Modeling in Mechanobiology 12.3 (2013), pp. 617–626. ISSN: 1617-7959, 1617-7940. DOI: 10.1007/s10237-012-0429-0.

[56] Grahame A. Busby et al. “Confined Compression of Collagen Hydrogels”. In: Journal of Biomechanics 46.4 (2013), pp. 837–840. ISSN: 00219290. DOI: 10.1016/j.jbiomech.2012.11.048.

[57] Steven H.V. Cornet et al. “Water release kinetics from soy protein gels and meat analogues as studied with confined compression”. In: Innovative Food Science & Emerging Technologies 66 (2020), p. 102528. DOI: 10.1016/j.ifset.2020.102528.

[58] Bryan A. Nerger et al. “Tuning Porosity of Macroporous Hydrogels Enables Rapid Rates of Stress Relaxation and Promotes Cell Expansion and Migration”. In: Proceedings of the National Academy of Sciences 121.45 (2024), e2410806121. ISSN: 0027-8424, 1091-6490. DOI: 10.1073/pnas.2410806121.

[59] Z. Ilke Kalcioglu et al. “From Macro-to Microscale Poroelastic Characterization of Polymeric Hydrogels via Indentation”. In: Soft Matter 8.12 (2012), p. 3393. ISSN: 1744-683X. DOI: 10.1039/c2sm06825g.

[60] Sureshkumar Kalyanam, Kathleen S. Toohey, and Michael F. Insana. “Modeling Biphasic Hydrogels under Spherical Indentation: Application to Soft Tissues”. In: Mechanics of Materials 161 (2021), p. 103987. ISSN: 01676636. DOI: 10.1016/j.mechmat.2021.103987.

[61] Manuel P. Kainz et al. “Poro-Viscoelastic Material Parameter Identification of Brain Tissue-Mimicking Hydrogels”. In: Frontiers in Bioengineering and Biotechnology 11 (2023), p. 1143304. ISSN: 2296-4185. DOI: 10.3389/fbioe.2023.1143304.

[62] Alexander Leibinger et al. “Soft Tissue Phantoms for Realistic Needle Insertion: a Comparative Study”. In: Annals of Biomedical Engineering 44.8 (2016), pp. 2442–2452. ISSN: 0090-6964. DOI: 10.1007/s10439-015-1523-0.

[63] Michele Terzano et al. “Fluid–Solid Interaction in the Rate-Dependent Failure of Brain Tissue and Biomimicking Gels”. In: Journal of the Mechanical Behavior of Biomedical Materials 119 (2021), p. 104530. ISSN: 17516161. DOI: 10.1016/j.jmbbm.2021.104530.

[64] Manuel P. Kainz et al. “Biointegration of Soft Tissue-Inspired Hydrogels on the Chorioallantoic Membrane: an Experimental Characterization”. In: Materials Today Bio 31 (2025), p. 101508. ISSN: 25900064. DOI: 10.1016/j.mtbio.2025.101508.

[65] Zhengchu Tan et al. “Composite Hydrogel: a High Fidelity Soft Tissue Mimic for Surgery”. In: Materials & Design 160 (2018), pp. 886–894. ISSN: 02641275. DOI: 10.1016/j.matdes.2018.10.018.

[66] Zhengchu Tan et al. “Cryogenic 3D Printing of Super Soft Hydrogels”. In: Scientific Reports 7.1 (2017), p. 16293. ISSN: 2045-2322. DOI: 10.1038/s41598-017-16668-9.

[67] Maurice A Biot. “General Theory of Three-dimensional Consolidation”. In: Journal of Applied Physics 12.2 (1941), pp. 155–164. ISSN: 0021-8979. DOI: 10.1063/1.1712886.

[68] Paul J. Flory and John Rehner. “Statistical Mechanics of Cross-Linked Polymer Networks II. Swelling”. In: The Journal of Chemical Physics 11.11 (1943), pp. 521–526. ISSN: 0021-9606, 1089-7690. DOI: 10.1063/1.1723792.

[69] Wei Hong et al. “A Theory of Coupled Diffusion and Large Deformation in Polymeric Gels”. In: Journal of the Mechanics and Physics of Solids 56.5 (2008), pp. 1779–1793. ISSN: 00225096. DOI: 10.1016/j.jmps.2007.11.010.

[70] Ray M. Bowen. “Incompressible porous media models by use of the theory of mixtures”. In: International Journal of Engineering Science 18.9 (1980), pp. 1129–1148. DOI: 10.1016/0020-7225(80)90114-7.

[71] Lallit Anand. “2014 Drucker Medal Paper: A Derivation of the Theory of Linear Poroelasticity From Chemoelasticity”. In: Journal of Applied Mechanics 82.11 (2015), p. 111005. ISSN: 0021-8936, 1528-9036. DOI: 10.1115/1.4031049.

[72] K. R. Rajagopal. “Diffusion through Polymeric Solids Undergoing Large Deformations”. In: Materials Science and Technology 19.9 (2003), pp. 1175–1180. ISSN: 02670836. DOI: 10.1179/026708303225004729.

[73] A.D. Drozdov et al. “Constitutive Equations for the Kinetics of Swelling of Hydrogels”. In: Mechanics of Materials 102 (2016), pp. 61–73. ISSN: 01676636. DOI: 10.1016/j.mechmat.2016.08.012.

[74] C. Truesdell and R. Toupin. “The classical field theories”. In: Principles of classical mechanics and field theory. Ed. by S. Flügge. Berlin, Heidelberg: Springer Berlin Heidelberg, 1960, pp. 226–858. ISBN: 978-3-642-45943-6. DOI: 10.1007/978-3-642-45943-6\_2.

[75] S. Majid Hassanizadeh and William G Gray. “General Conservation Equations for Multi-Phase Systems: 1. Averaging Procedure”. In: Advances in Water Resources 2 (1979), pp. 131–144. ISSN: 03091708. DOI: 10.1016/0309-1708(79)90025-3.

[76] Gerard A. Ateshian. “Mixture Theory for Modeling Biological Tissues: illustrations from Articular Cartilage”. In: Biomechanics: Trends in Modeling and Simulation. Ed. by Gerhard A. Holzapfel and Ray W. Ogden. Vol. 20. Cham: Springer International Publishing, 2017, pp. 1–51. DOI: 10.1007/978-3-319-41475-1\_1.

[77] Christopher W. MacMinn, Eric R. Dufresne, and John S. Wettlaufer. “Large Deformations of a Soft Porous Material”. In: Physical Review Applied 5.4 (2016), p. 044020. ISSN: 2331-7019. DOI: 10.1103/PhysRevApplied.5.044020.

[78] Wolfgang Ehlers and G Eipper. “Finite Elastic Deformations in Liquid-Saturated and Empty Porous Solids”. In: Porous Media: Theory and Experiments. Vol. 34. Dordrecht: Springer Netherlands, 1999, pp. 179–191.

[79] Gerhard Sommer et al. “Multiaxial Mechanical Properties and Constitutive Modeling of Human Adipose Tissue: a Basis for Preoperative Simulations in Plastic and Reconstructive Surgery”. In: Acta Biomaterialia 9.11 (2013), pp. 9036–9048. ISSN: 1742-7061. DOI: 10.1016/j.actbio.2013.06.011.

[80] Henry W Haslach et al. “Solid-Extracellular Fluid Interaction and Damage in the Mechanical Response of Rat Brain Tissue under Confined Compression”. In: Journal of the Mechanical Behavior of Biomedical Materials 29 (2014), pp. 138–150. DOI: 10.1016/j.jmbbm.2013.08.027.

[81] D.E. Liu et al. “Equilibrium Water and Solute Uptake in Silicone Hydrogels”. In: Acta Biomaterialia 18 (2015), pp. 112–117. ISSN: 17427061. DOI: 10.1016/j.actbio.2015.02.019.

[82] Caroline A Schneider, Wayne S Rasband, and Kevin W Eliceiri. “NIH Image to ImageJ: 25 Years of Image Analysis”. In: Nature Methods 9.7 (2012), pp. 671–675. ISSN: 1548-7091, 1548-7105. DOI: 10.1038/nmeth.2089.

[83] R. Hilfer. “Review on Scale Dependent Characterization of the Microstructure of Porous Media”. In: Transport in Porous Media 46.2-3 (2002), pp. 373–390. ISSN: 0169-3913, 1573-1634. DOI: 10.1023/A:1015014302642.

[84] Yan Tang et al. “Behaviors of Water Molecules in Polyvinyl Alcohol Gel amid Stretch and Temperature Changes: a Molecular Dynamics Study”. In: Materials Today Communications 33 (2022), p. 104834. ISSN: 23524928. DOI: 10.1016/j.mtcomm.2022.104834.

[85] Georgia Kaklamani et al. “Mechanical Properties of Alginate Hydrogels Manufactured Using External Gelation”. In: Journal of the Mechanical Behavior of Biomedical Materials 36 (2014), pp. 135–142. ISSN: 17516161. DOI: 10.1016/j.jmbbm.2014.04.013.

[86] Silvia Todros et al. “Time-Dependent Mechanical Behavior of Partially Oxidized Polyvinyl Alcohol Hydrogels for Tissue Engineering”. In: Journal of the Mechanical Behavior of Biomedical Materials 125.July 2021 (2022), p. 104966. ISSN: 17516161. DOI: 10.1016/j.jmbbm.2021.104966.

[87] Yousef Javanmardi et al. “Quantifying Cell-Generated Forces: Poisson’s Ratio Matters”. In: Communications Physics 4.1 (2021), p. 237. ISSN: 2399-3650. DOI: 10.1038/s42005-021-00740-y.

[88] Nathan R. Richbourg, Manuel K. Rausch, and Nicholas A. Peppas. “Cross-Evaluation of Stiffness Measurement Methods for Hydrogels”. In: Polymer 258 (2022), p. 125316. ISSN: 00323861. DOI: 10.1016/j.polymer.2022.125316.

[89] Ariell Marie Smith et al. “Facile Determination of the Poisson’s Ratio and Young’s Modulus of Polyacrylamide Gels and Polydimethylsiloxane”. In: ACS Applied Polymer Materials 6.4 (2024), pp. 2405–2416. ISSN: 2637-6105, 2637-6105. DOI: 10.1021/acsapm.3c03154.

[90] Shawn A Chester and Lallit Anand. “A Coupled Theory of Fluid Permeation and Large Deformations for Elastomeric Materials”. In: Journal of the Mechanics and Physics of Solids 58.11 (2010), pp. 1879–1906. ISSN: 00225096. DOI: 10.1016/j.jmps.2010.07.020.

[91] S. Baek and T.J. Pence. “Inhomogeneous Deformation of Elastomer Gels in Equilibrium under Saturated and Unsaturated Conditions”. In: Journal of the Mechanics and Physics of Solids 59.3 (2011), pp. 561–582. ISSN: 00225096. DOI: 10.1016/j.jmps.2010.12.013.

[92] Alessandro Lucantonio, P. Nardinocchi, and L. Teresi. “Transient Analysis of Swelling-Induced Large Deformations in Polymer Gels”. In: Journal of the Mechanics and Physics of Solids 61.1 (2013), pp. 205–218. ISSN: 00225096. DOI: 10.1016/j.jmps.2012.07.010.

[93] Gerard A. Ateshian and Jeffrey A. Weiss. “Anisotropic Hydraulic Permeability Under Finite Deformation”. In: Journal of Biomechanical Engineering 132.11 (2010), pp. 1–7. ISSN: 0148-0731. DOI: 10.1115/1.4002588.

[94] Mattia Bacca, Omar A. Saleh, and Robert M. McMeeking. “Contraction of Polymer Gels Created by the Activity of Molecular Motors”. In: Soft Matter 15.22 (2019), pp. 4467–4475. ISSN: 1744-683X, 1744-6848. DOI: 10.1039/C8SM02598C.

[95] Matteo Ferraresso et al. “Energetics of Cytoskeletal Gel Contraction”. In: Soft Matter 19.29 (2023), pp. 5430– 5442. ISSN: 1744-683X, 1744-6848. DOI: 10.1039/D2SM01557A.

[96] Zelai Xu, Pengtao Yue, and James J. Feng. “A Theory of Hydrogel Mechanics That Couples Swelling and External Flow”. In: Soft Matter 20.27 (2024), pp. 5389–5406. ISSN: 1744-683X, 1744-6848. DOI: 10.1039/D4SM00424H.

[97] W. M. Lai, J. S. Hou, and V. C. Mow. “A Triphasic Theory for the Swelling and Deformation Behaviors of Articular Cartilage”. In: Journal of Biomechanical Engineering 113.3 (1991), pp. 245–258. ISSN: 0148-0731, 1528-8951. DOI: 10.1115/1.2894880.

[98] Gerard A. Ateshian, Steve Maas, and Jeffrey A. Weiss. “Multiphasic Finite Element Framework for Modeling Hydrated Mixtures With Multiple Neutral and Charged Solutes”. In: Journal of Biomechanical Engineering 135.11 (2013), p. 111001. ISSN: 0148-0731, 1528-8951. DOI: 10.1115/1.4024823.

[99] Qi-Ming Wang et al. “Separating Viscoelasticity and Poroelasticity of Gels with Different Length and Time Scales”. In: Acta Mechanica Sinica 30.1 (2014), pp. 20–27. ISSN: 0567-7718. DOI: 10.1007/s10409-014-0015-z.

[100] Lijun Su et al. “Distinguishing Poroelasticity and Viscoelasticity of Brain Tissue with Time Scale”. In: Acta Biomaterialia 155 (2023), pp. 423–435. ISSN: 17427061. DOI: 10.1016/j.actbio.2022.11.009.

[101] Stéphane Cuenot et al. “Poroelastic and Viscoelastic Properties of Soft Materials Determined from AFM Force Relaxation and Force-Distance Curves”. In: Journal of the Mechanical Behavior of Biomedical Materials 163 (2025), p. 106865. ISSN: 17516161. DOI: 10.1016/j.jmbbm.2024.106865.

[102] Yuhang Hu and Zhigang Suo. “Viscoelasticity and Poroelasticity in Elastomeric Gels”. In: Acta Mechanica Solida Sinica 25.5 (2012), pp. 441–458. DOI: 10.1016/S0894-9166(12)60039-1.

[103] Imanda Jayawardena et al. “Evaluation of Techniques Used for Visualisation of Hydrogel Morphology and Determination of Pore Size Distributions”. In: Materials Advances 4.2 (2023), pp. 669–682. ISSN: 2633-5409. DOI: 10.1039/D2MA00932C.

[104] Dimitra Katrantzi et al. “Unveiling the Structure of Protein-Based Hydrogels by Overcoming Cryo-SEM Sample Preparation Challenges”. In: Faraday Discussions 260 (2025), pp. 55–81. ISSN: 1359-6640, 1364-5498. DOI: 10.1039/D4FD00204K.

[105] Shan Jiang, Sha Liu, and Wenhao Feng. “PVA Hydrogel Properties for Biomedical Application”. In: Journal of the Mechanical Behavior of Biomedical Materials 4.7 (2011), pp. 1228–1233. ISSN: 17516161. DOI: 10.1016/j.jmbbm.2011.04.005.

[106] Shirsha Bose, Majid Akbarzadeh Khorshidi, and Caitríona Lally. “Tailoring the Mechanical Properties of Macro-Porous PVA Hydrogels for Biomedical Applications”. In: Journal of the Mechanical Behavior of Biomedical Materials 161 (2025), p. 106787. ISSN: 17516161. DOI: 10.1016/j.jmbbm.2024.106787.

[107] Ruigang Zhou et al. “Multifunctional Hydrogel Based on Polyvinyl Alcohol/Chitosan/Metal Polyphenols for Facilitating Acute and Infected Wound Healing”. In: Materials Today Bio 29 (2024), p. 101315. ISSN: 25900064. DOI: 10.1016/j.mtbio.2024.101315.

[108] Vikram Singh Raghuwanshi and Gil Garnier. “Characterisation of Hydrogels: linking the Nano to the Microscale”. In: Advances in Colloid and Interface Science 274 (2019), p. 102044. ISSN: 00018686. DOI: 10.1016/j.cis.2019.102044.

[109] Jeanne Aigoin et al. “Comparative Analysis of Electron Microscopy Techniques for Hydrogel Microarchitecture Characterization: SEM, Cryo-SEM, ESEM, and TEM”. In: ACS Omega 10.15 (2025), pp. 14687–14698. ISSN: 2470-1343, 2470-1343. DOI: 10.1021/acsomega.4c08096.

[110] Abner Velazco et al. “Reduction of SEM Charging Artefacts in Native Cryogenic Biological Samples”. In: Nature Communications 16.1 (2025), p. 5204. ISSN: 2041-1723. DOI: 10.1038/s41467-025-60545-3.

