## Supplementary Material for "Effective porosity and fluid flow in macroporous ultrasoft hydrogels: An experimental characterization"

**List of contents**

1 Surface porosity evaluation via ImageJ thresholding (supplementing Sec. 3.4.1 Image processing)

2 Total fluid fraction and effective porosity depending on the polymer concentrations (supplementing Sec. 4.3 Incremental consolidation: free-water fraction and point of compaction)

3 References

### 1 Surface porosity evaluation via ImageJ thresholding (supplementing Sec. 3.4.1 Image processing)

To assess the surface porosity of cryo-SEM images of hydrogel samples, a standardized image processing workflow was established using the open-source software *ImageJ* (Schneider et al., 2012). We closely followed the validated process described by Hojat et al., 2022. First, an original micrograph was opened and a rectangular region of interest (ROI) was selected to exclude the image label and scale bar. This ROI was extracted using *Image > Scale* to ensure consistent area comparison across samples. Original images, including the image labels and scale bars, are shown in Fig.1(a),(c),(e). After separating the image and the label, a grayscale threshold was applied (*Image > Adjust > Threshold*) to enhance contrast between the polymer network in the foreground and the darker, frozen water in the background. Typical histograms revealed two distinct peaks in all our data, since the darker background was clearly distinguishable from the lighter polymer matrix in the front, thus enabling robust segmentation of the hydrogel matrix versus background. Different threshold styles (e.g., 'Red', 'B&W', 'Over/Under') were systematically tested, but did not significantly alter the final values. The 'default' threshold style was therefore used for all analyses. Binary images were generated (*Process > Binary > Make Binary*) and further post-processed to eliminate artifacts caused by sample preparation (e.g., ice crystals). A summary of the available thresholds and the resulting binary images is shown in Fig.2.

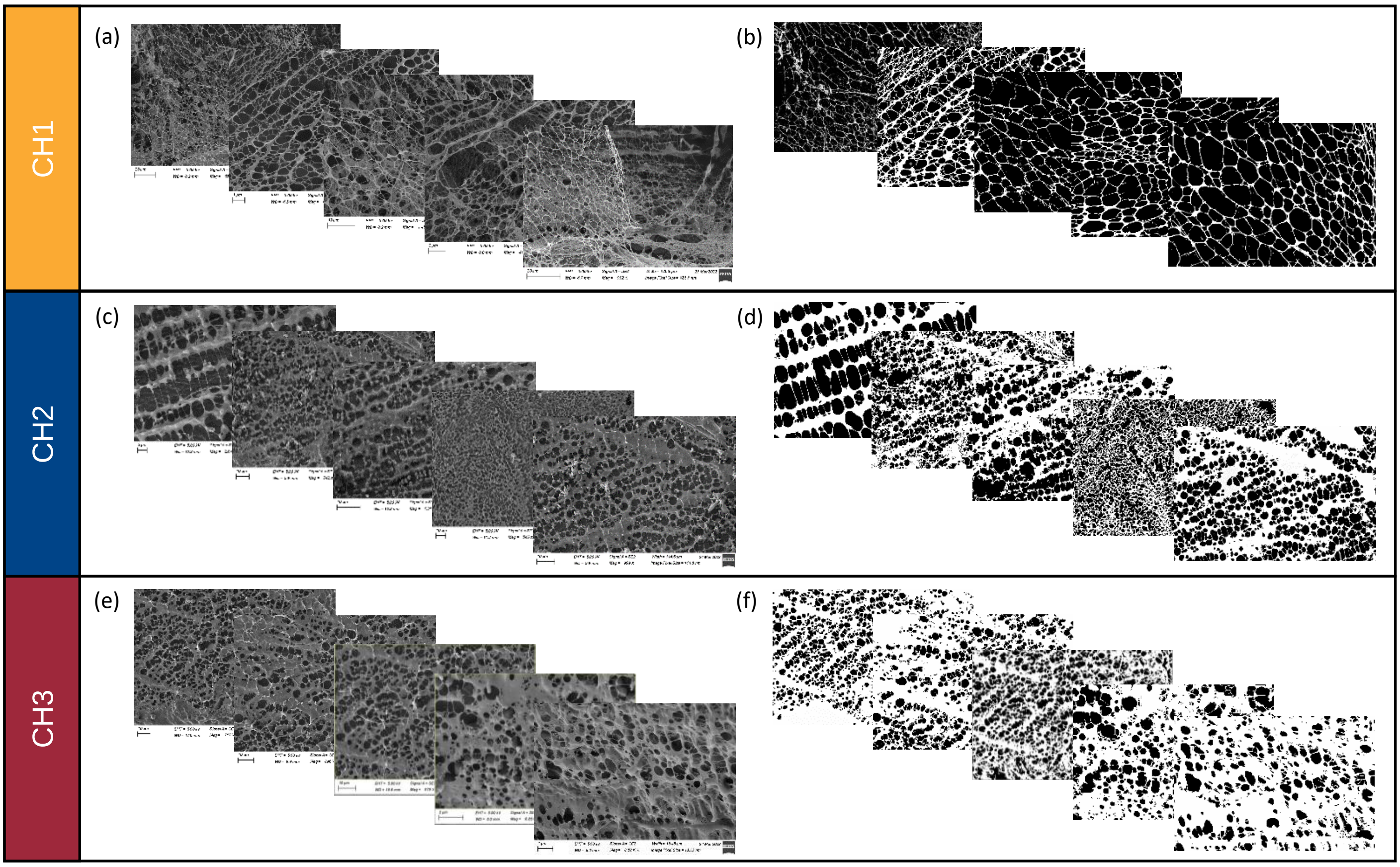

Figure 1. Cryo-SEM image analyses and binarized images as used for image processing.

An artefact correction was applied. Without this correction, artifacts with similar grayscale values to the hydrogel could falsely be included in the solid phase, resulting in overestimation of the solid/matrix area. The steps involved are shown in Fig.3 (a)-(f). Following the correction, the ratio of pore area and total area was obtained using *Analyze > Analyze Particles*. This step returns a visualization of the pores (see Fig.3(g)) as well as a table with the corresponding metrics. The "Area %" output, defined as the ratio of pore space to total ROI area, was used as the surface porosity metric.

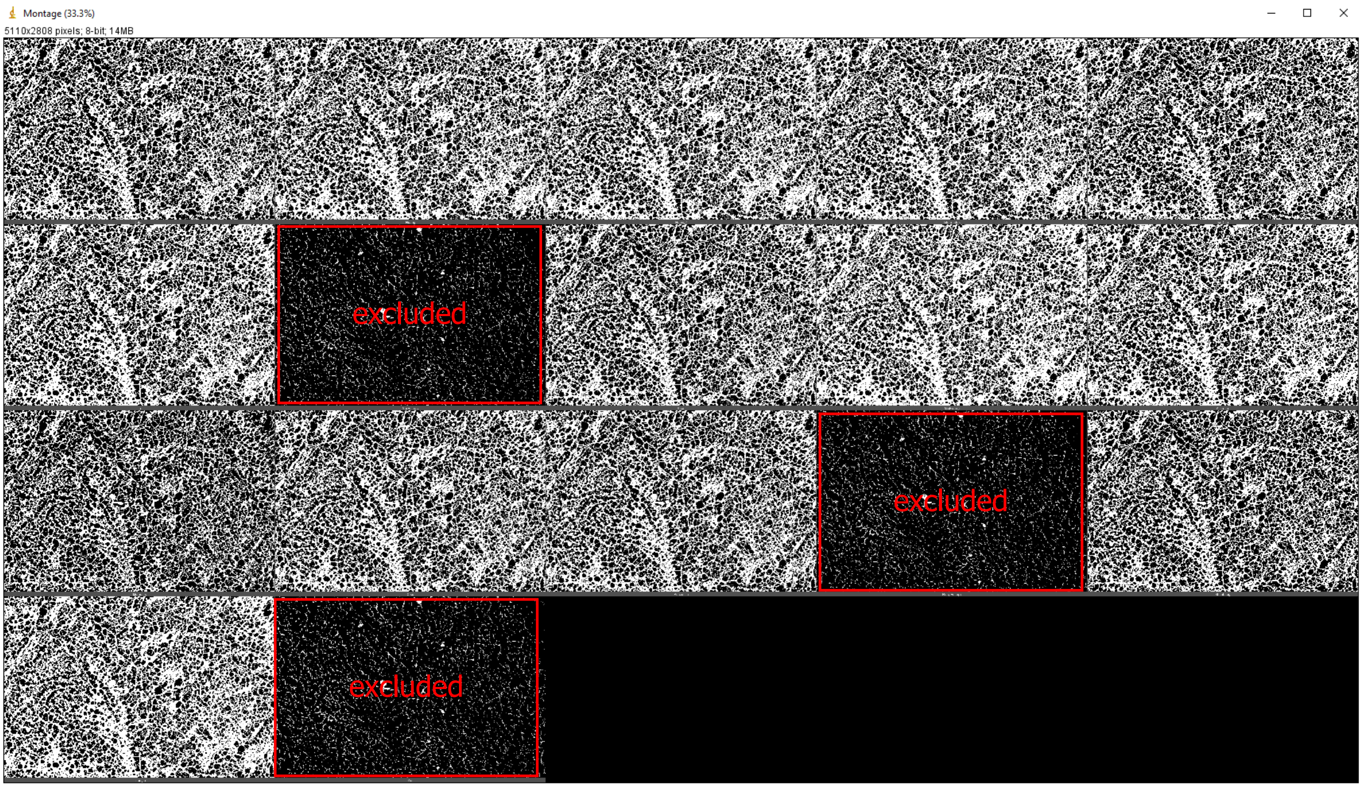


Figure 2: Threshold variation to test the influence of surface porosity ('%Area') after extraction of the frame and before transformation to a binary image. Example: Composite hydrogel CH2.

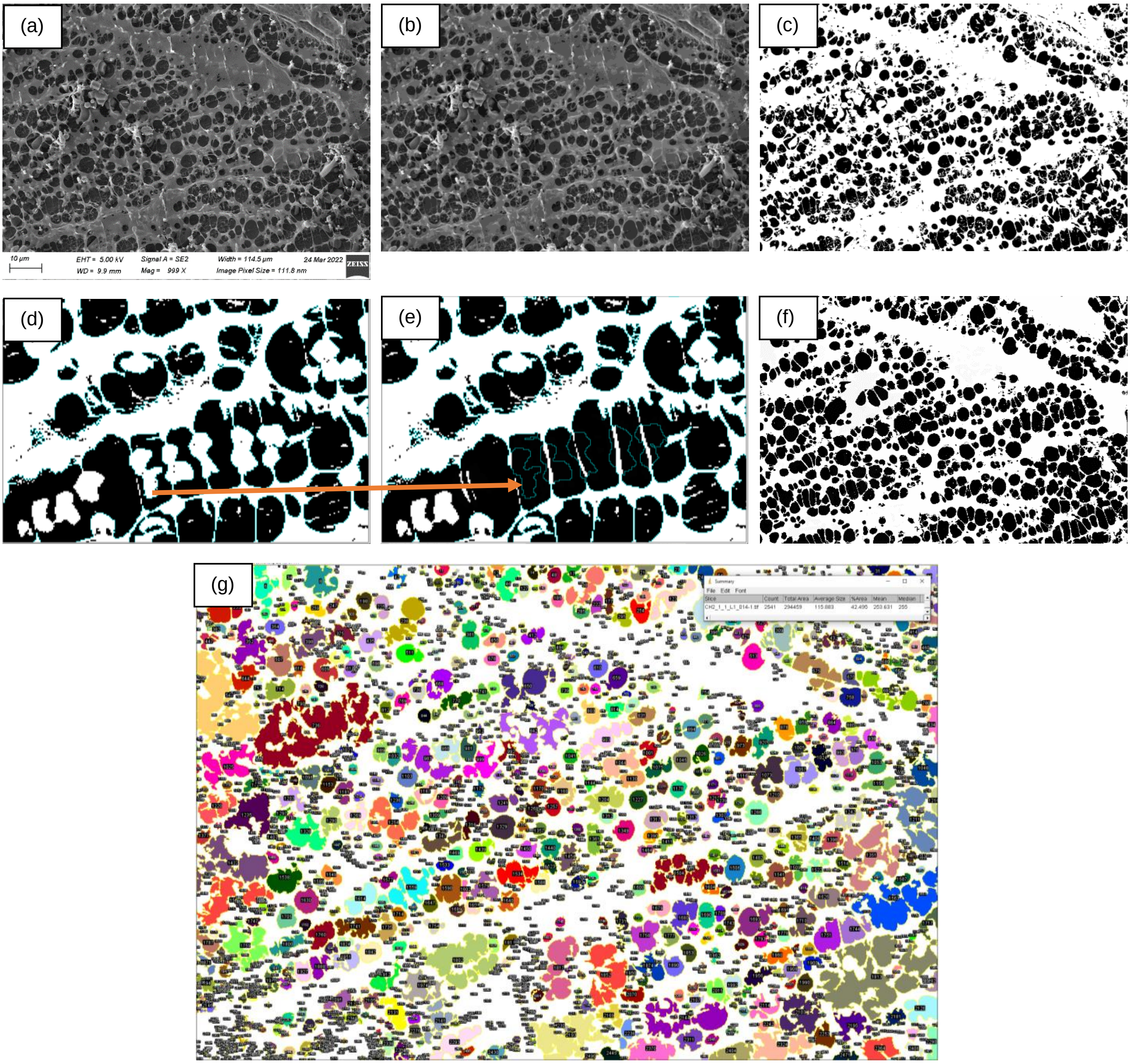


Figure 3: Example of all steps involved in the workflow to derive the surface porosity from cryo-SEM images. Example: Composite hydrogel CH2.

### 2 Total fluid fraction and effective porosity depending on the polymer concentrations (supplementing Sec. 4.3 Incremental consolidation: free-water fraction and point of compaction)

Higher polymer concentrations consistently lead to lower water content across all hydrogel formulations. Fig. 4 (a) shows an approximately linear decrease in total fluid fraction with increasing total polymer concentration. Fig. 4 (b),(c),(d) show that the effective porosity drops systematically not only with total polymer content but also when polyvinyl alcohol (PVA) and Phytagel (PHY) concentrations are considered individually. Together, these trends illustrate that the mobile-water fraction is directly tunable through formulation parameters.

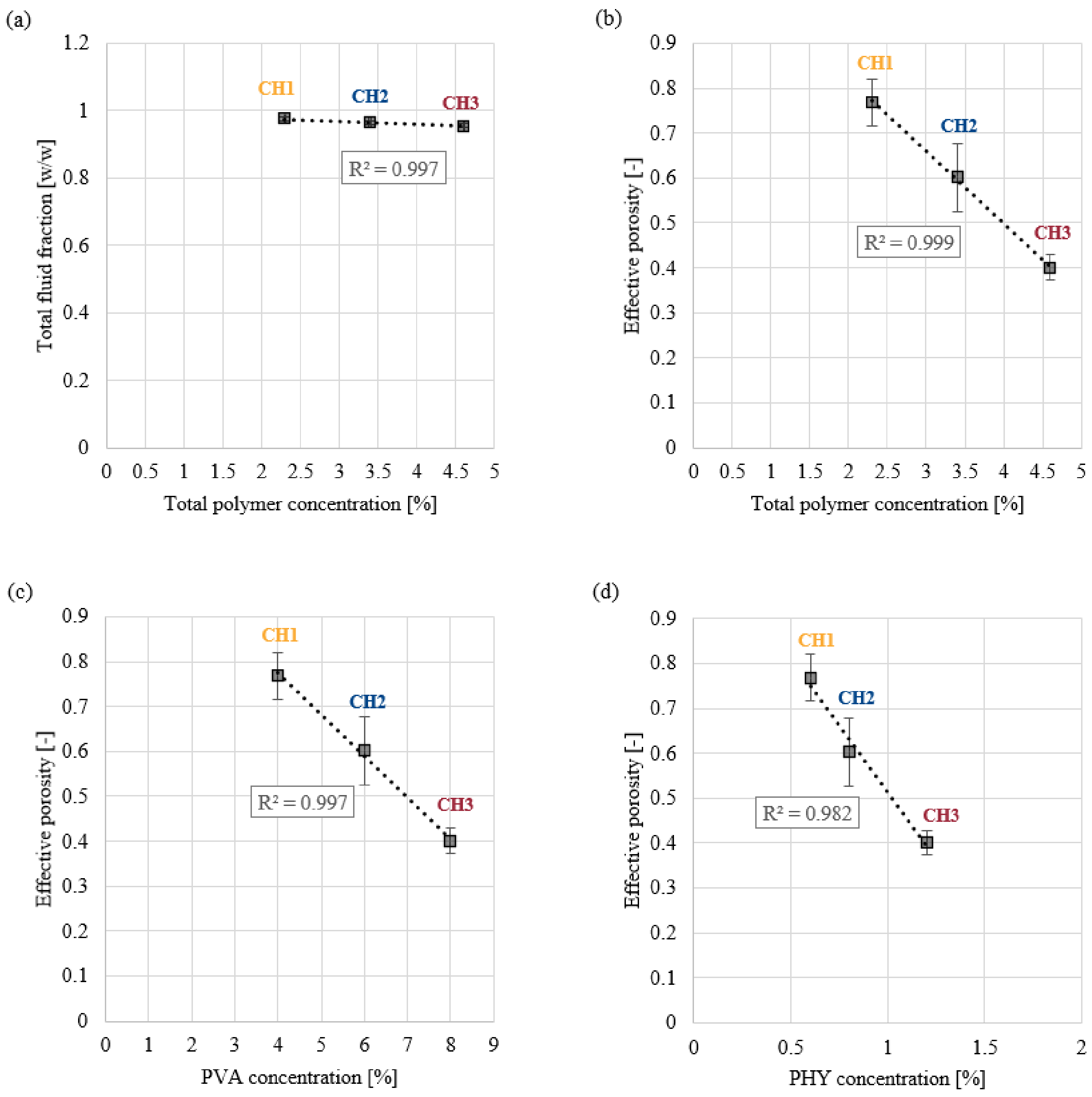


Figure 4. Fluid fraction and effective porosity depending on the polymer content. (a) Total fluid fraction after dehydration over total polymer concentration. Effective porosity (amount of free water) depending on (b) total polymer concentration, (c) concentration of PVA and (d) concentration of PHY.

### 3 References


Hojat, N., Gentile, P., Ferreira, A.M., Šiller, L. Automatic pore size measurements from scanning electron microscopy images of porous scaffolds. J. Porous Mater. 30, 93–101 (2023). doi:10.1007/s10934-022-01309-y.

Schneider, C.A., Rasband, W.S., Eliceiri, K.W. NIH Image to ImageJ: 25 years of image analysis. Nat. Methods 9, 671–675 (2012). doi:10.1038/nmeth.2089.
